# Genetically Encoded Photodecaging of Asparagine Enables Protein Function Control and Time-Resolved N-Glycosylation

**DOI:** 10.64898/2026.11.24.768507

**Authors:** Xianrui Zhang, Zhouchang Xu, Zhiyuan Zhang, Peng R. Chen

**Author notes:** Correspondence to: Peng R. Chen. These authors contributed equally to this work (Xianrui Zhang, Zhouchang Xu, Zhiyuan Zhang).

## Abstract

Asparagine contributes to protein function through its polar carboxamide side chain and serves as the acceptor residue for N-linked glycosylation, yet temporally controlling these roles directly through the side-chain carboxamide remains challenging. Here, we report the genetic encoding of *N*^4^-(2-nitrobenzyl)-L-asparagine (ONBN), a photocaged asparagine that regenerates the native side-chain carboxamide upon 365 nm irradiation. Engineered aminoacyl-tRNA synthetase/tRNA pairs enabled site-specific ONBN incorporation into proteins in *Escherichia coli* and mammalian cells. Photodecaging restored *Renilla* luciferase activity and activated TRAF6 polyubiquitination mediated by UBE2N in living cells. Beyond active-site residues, we further developed CAGE-prox^N^, using computationally guided proximal caging to optically activate firefly luciferase. Finally, ONBN incorporation at the acceptor position of an N-glycosylation sequon enabled light-triggered initiation and time-resolved monitoring of post-translational glycan installation on an endoplasmic reticulum-targeted reporter. Variation of the −2 flanking residue revealed sequence-dependent differences in apparent glycosylation kinetics. This work extends genetically encoded photocontrol to the asparagine carboxamide and provides a platform for investigating asparagine-dependent protein functions and the temporal regulation of N-glycosylation in living cells.

## Introduction

Site-specific manipulation of amino acid side chains provides a precise approach for investigating and controlling protein function^1,2^. In particular, genetic code expansion (GCE) combined with bioorthogonal decaging reactions enables a selected amino acid side chain to be transiently masked and subsequently restored on demand^3,4^. In this strategy, a caged unnatural amino acid (UAA) is genetically incorporated at a defined site within a protein of interest, where it suppresses protein activity or molecular interactions until removal of the cage by light or a chemical trigger^5–8^. Regeneration of the natural amino acid restores the native protein sequence and function, enabling gain-of-function manipulation with high spatiotemporal precision. These approaches have been applied to the side chains of Ser^9^, Tyr^10^, Cys^11^, Lys^12^, His^13^, Glu^14–16^, Asp^17^, Trp^18^ and Phe^19^, enabling the regulation of enzymatic activity, molecular recognition, protein–protein interactions and cellular signaling^20–24^.

Despite these advances, among the functional groups found in the 20 natural amino acids, the carboxamide side chains of asparagine (Asn) and glutamine (Gln) remain conspicuously absent from the genetically encoded bioorthogonal decaging toolkit. At physiological pH, these side chains are neutral but highly polar: their carbonyl oxygen and amide N-H groups function as hydrogen-bond acceptors and donors, respectively^25^. This combination enables directional interactions that organize protein hydrogen-bond networks, stabilize buried polar architectures^26^, position substrates and catalytic residues, and contribute to molecular recognition without introducing formal charge (Fig. 1a)^27–29^. Beyond these structural and functional roles, Asn serves as the acceptor residue for protein N-glycosylation, which commonly occurs within an Asn-X-Ser/Thr sequon^30^ and is strongly influenced by the surrounding sequence and substrate structure^31–33^. These dual roles make Asn an especially compelling target for GCE-based bioorthogonal decaging, which would extend site-specific control to carboxamide-mediated protein functions while providing a unique means to temporally regulate and interrogate N-glycosylation.

**Figure 1.**
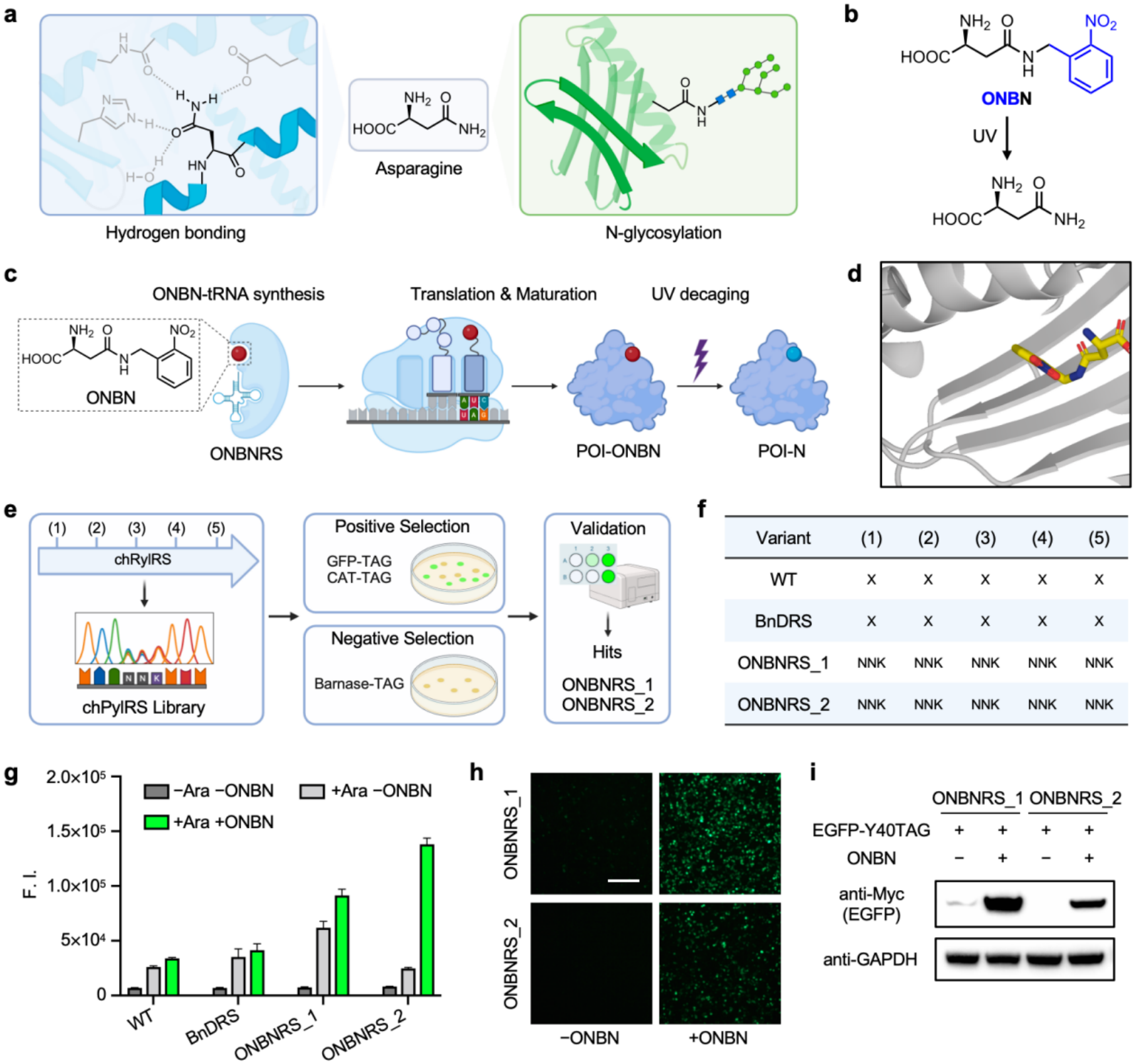
Development and genetic encoding of a photocaged asparagine. **(a)** Hydrogen-bonding and N-glycosylation functions of asparagine. **(b)** Photodecaging of ONBN to regenerate asparagine. **(c)** ONBNRS-mediated tRNA aminoacylation, amber suppression, protein maturation and photodecaging of the protein of interest (POI). **(d)** Predicted ONBN–AMP-binding pocket of BnDRS. **(e)** Focused NNK library construction, positive selection with GFPuv-D190TAG and chloramphenicol acetyltransferase (CAT)-D112TAG reporters, barnase-TAG negative selection and hit validation. **(f)** Schematic of the five library positions for reference chPylRS (WT), BnDRS, ONBNRS_1 and ONBNRS_2. N and NNK denote masked positions and do not report individual residue identities. **(g)** sfGFP-D190TAG fluorescence in *E. coli* DH10B with the indicated L-arabinose (Ara) and ONBN conditions. Data are mean ± SD (*n* = 4). F.I., fluorescence intensity. **(h)** HEK293T cells expressing EGFP-Y40TAG with ONBNRS_1 or ONBNRS_2, with or without ONBN. Scale bar, 200 μm. (i) Corresponding anti-Myc immunoblot of EGFP, with GAPDH as the loading control. Immunoblots are representative of three independent experiments.

Here, we report the genetic encoding of *N*^4^-(2-nitrobenzyl)-L-asparagine (ONBN), a photocaged Asn derivative that enables bioorthogonal, light-triggered regeneration of the natural carboxamide side chain. Through aminoacyl-tRNA synthetase engineering, we achieved site-specific incorporation of ONBN into proteins in *E. coli* and mammalian cells. We applied ONBN decaging to optically activate *Renilla* luciferase (RLuc) and UBE2N-mediated TRAF6 polyubiquitination, targeting Asn residues implicated in hydrogen bonding and organization of the ubiquitin-transfer active site, respectively. We further developed CAGE-prox^N^, a computationally guided proximal-caging strategy, and demonstrated its utility for optical control of firefly luciferase (FLuc). Beyond protein activation, we incorporated ONBN into an N-glycosylation sequon of endoplasmic reticulum (ER)-targeted enhanced green fluorescent protein (EGFP) to initiate post-translational N-glycosylation by light in living cells. Time-resolved analysis of reporters bearing different residues at the −2 position revealed sequence-dependent glycosylation kinetics. Together, this work extends site-specific gain-of-function control to the Asn carboxamide and provides a platform for interrogating Asn-mediated protein functions and temporally regulating N-glycosylation.

## Results and discussion

### Development and genetic encoding of photocaged asparagine

To enable light-controlled manipulation of asparagine (Asn) side chains, we designed a photocaged Asn derivative, ONBN, in which the side-chain amide nitrogen is modified with an *o*-nitrobenzyl group (Fig. 1b). This design exploits the photolability of *o*-nitrobenzyl protecting groups^15,17,34^ to regenerate the Asn carboxamide upon 365 nm irradiation. We synthesized and characterized ONBN, then evaluated its photodecaging kinetics at the small-molecule level under physiologically relevant conditions. These analyses confirmed the light-triggered release of Asn from ONBN (Fig. S1).

We next sought to develop an aminoacyl-tRNA synthetase (aaRS)/tRNA pair capable of incorporating ONBN into proteins through amber codon suppression^35^ (Fig. 1c). Guided by our previously established computational workflow for noncanonical amino acid–synthetase engineering^16^, we selected BnDRS, which recognizes aspartic acid β-benzyl ester (BnD)^36^, as the parental chimeric PylRS scaffold^37,38^. AlphaFold 3^39^ was used to predict the BnDRS complex with the ONBN adenylate intermediate (ONBN–AMP; Fig. 1d; Fig. S2). Five pocket positions were prioritized for focused NNK mutagenesis. Positive and negative selection, followed by reporter-based validation, yielded two synthetase variants, designated ONBNRS_1 and ONBNRS_2 (Fig. 1e,f; Fig. S3).

To evaluate their performance in bacterial and mammalian cells^12^, we introduced each synthetase/tRNA pair into pSUPAR for bacterial expression and pRS (Fig. S3) for mammalian expression. In *Escherichia coli*, a superfolder green fluorescent protein (sfGFP) reporter carrying D190TAG showed increased fluorescence upon ONBN supplementation with either variant, and both produced substantially higher fluorescence than reference chPylRS or parental BnDRS under ONBN-supplemented conditions (Fig. 1g; Figs. S3, S5). ONBNRS_2 yielded a higher mean fluorescence signal than ONBNRS_1 and exhibited lower background fluorescence in the absence of ONBN. In HEK293T cells, we evaluated the variants using an EGFP-Y40TAG reporter. Fluorescence microscopy and immunoblotting showed increased EGFP expression upon ONBN supplementation with both variants (Fig. 1h,i; Fig. S6). ONBNRS_1 produced a stronger EGFP signal under the conditions tested. Together, chemical synthesis, computationally guided directed evolution, and cellular reporter assays yielded ONBN-responsive aaRS/tRNA systems functional in both bacterial and mammalian cells, providing a foundation for subsequent studies of Asn-dependent biological processes.

### ONBN photodecaging in model proteins and functional validation in living cells

Having established aaRS/tRNA systems supporting ONBN incorporation, we next examined protein incorporation and photodecaging at the molecular level. Bacterial sfGFP-D190ONBN prepared with ONBNRS_2 was characterized by intact-protein UPLC–MS and tryptic peptide LC–MS/MS (Figs. S7, S8). Myc-tagged EGFP-Y40ONBN was expressed with ONBNRS_1 in HEK293T cells. HEK293T cells were either left unirradiated or irradiated for 2 min. Following cell harvest and lysis, EGFP was enriched and analyzed by intact-protein UPLC–MS. The calculated and observed masses were 28,103 and 28,102 Da for EGFP-Y40ONBN and 27,968 and 27,974 Da for EGFP-Y40N, respectively. The observed mass shift was consistent with removal of the ONBN protecting group within the estimated intact-mass uncertainty (Fig. 2a; Fig. S9). Cellular compatibility across ONBN concentrations and incubation times was assessed by MTT (Fig. S10).

**Figure 2.**
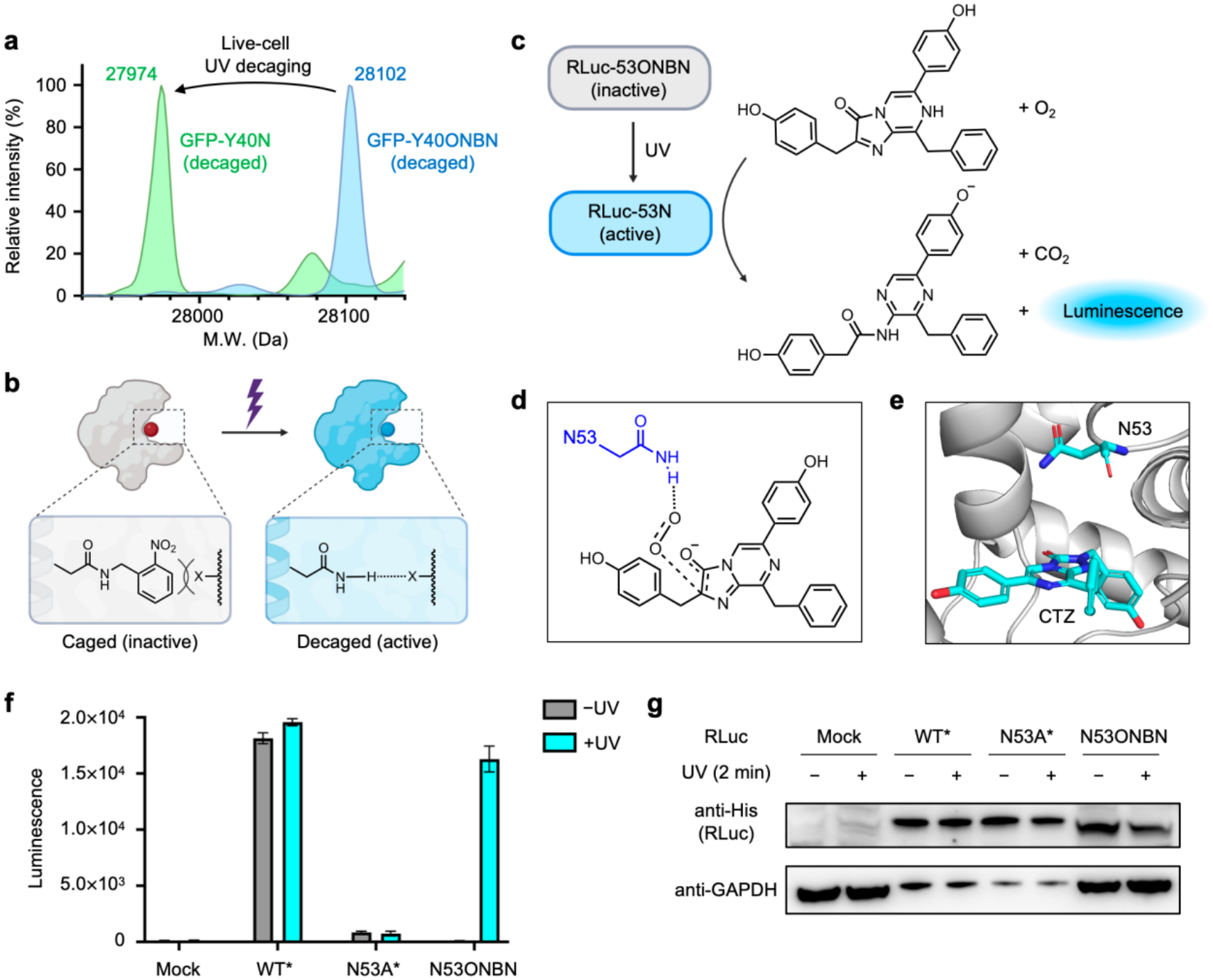
Optical control of model proteins through ONBN decaging. **(a)** Deconvoluted intact-protein mass spectra of Myc-tagged EGFP-Y40ONBN expressed with ONBNRS_1 before and after 2 min irradiation of intact cells. Each trace was independently normalized to a maximum intensity of 1; calculated and observed masses are given in Figure S9. **(b)** Schematic of protein photoactivation through decaging of ONBN and restoration of an Asn-mediated hydrogen-bonding interaction. **(c)** Photodecaging of RLuc-N53ONBN to RLuc-N53N and the RLuc-catalyzed oxidation of coelenterazine to generate coelenteramide, CO_2_, and light. RLuc, *Renilla* luciferase. **(d)** Hydrogen-bond-mediated positioning of molecular oxygen by N53 in the proposed RLuc catalytic mechanism. **(e)** Structural environment of N53 and coelenterazine (CTZ) in the RLuc active site. **(f)** Luminescence of Mock, WT RLuc, N53A and N53ONBN lysates after dark treatment or 2 min live-cell UV irradiation and immediate harvest. Lysate input was adjusted by immunoblotting to give comparable RLuc abundance. Bars show mean ± SD (*n* = 3). Asterisks mark the expression-adjusted WT and N53A lysates. **(g)** Anti-His immunoblot showing RLuc abundance after lysate-input adjustment, with GAPDH detected in the corresponding samples. Immunoblots are representative of three independent experiments.

We next investigated whether ONBN photodecaging could enable optical activation of a functional protein by restoring an Asn side chain involved in catalysis (Fig. 2b). *Renilla* luciferase (RLuc) catalyzes the oxygen-dependent oxidative decarboxylation of coelenterazine to generate coelenteramide, carbon dioxide, and luminescence^28,40^ (Fig. 2c). Mechanistic studies of *Renilla*-type luciferases identify N53 and W121 as hydrogen-bonding residues that position molecular oxygen for coelenterazine oxidation^28^ (Fig. 2d,e). We expressed RLuc-N53ONBN with ONBNRS_1 in HEK293T cells and compared luminescence after dark treatment or 2 min irradiation of intact cells followed by immediate harvest, alongside mock-transfected cells, WT RLuc, and RLuc-N53A (Fig. 2f,g). After irradiation and cell harvest, immunoblotting was used to adjust lysate amounts so that WT, N53A and N53ONBN samples contained comparable RLuc abundance across the −UV and +UV conditions (unadjusted activity and expression, Fig. S11). Without irradiation, RLuc-N53ONBN exhibited minimal luminescence. Following irradiation, RLuc-N53ONBN luminescence was 171.5 times that of the nonirradiated control (*n* = 3). These results demonstrate light-dependent recovery of RLuc activity following irradiation of living cells and support the use of genetically encoded ONBN for optical control of Asn-dependent protein function.

### Optical control of UBE2N-mediated TRAF6 polyubiquitination

Asparagine side chains can function as both hydrogen-bond donors and acceptors, contributing to the organization of catalytic microenvironments. E2 ubiquitin-conjugating enzymes receive activated ubiquitin (Ub) from E1 and cooperate with E3 ligases to transfer Ub to acceptor lysine residues^41^. In many E2 ubiquitin-conjugating enzymes, a conserved Asn resides within the HPN motif (Fig. 3a)^29^. In UBE2N (Ubc13), an E2 enzyme that catalyzes Lys63-linked ubiquitin chain formation in NF-κB signaling and DNA-damage repair^42^, this conserved Asn corresponds to N79. In the proposed oxyanion-stabilization model, the side-chain amide of N79 donates a hydrogen bond to the negatively charged C-terminal carbonyl oxygen of donor Ub during ubiquitin transfer^43–46^. We therefore selected N79 for ONBN incorporation, reasoning that photocaging could disrupt the interactions required for catalysis, whereas photodecaging would restore the native Asn side chain and enable enzyme activation. TRAF6 was used as a ubiquitination substrate to evaluate this optical control in living cells (Fig. 3b). Structural views locate N79 within the E2–cofactor–Ub complex (Fig. 3c) and illustrate its proximity to the donor-Ub C terminus in the active site (Fig. 3d)^47,48^. Such optical control could allow ubiquitination to be initiated at defined times and locations, providing a means to investigate how its timing and spatial organization influence downstream cellular responses.

**Figure 3.**
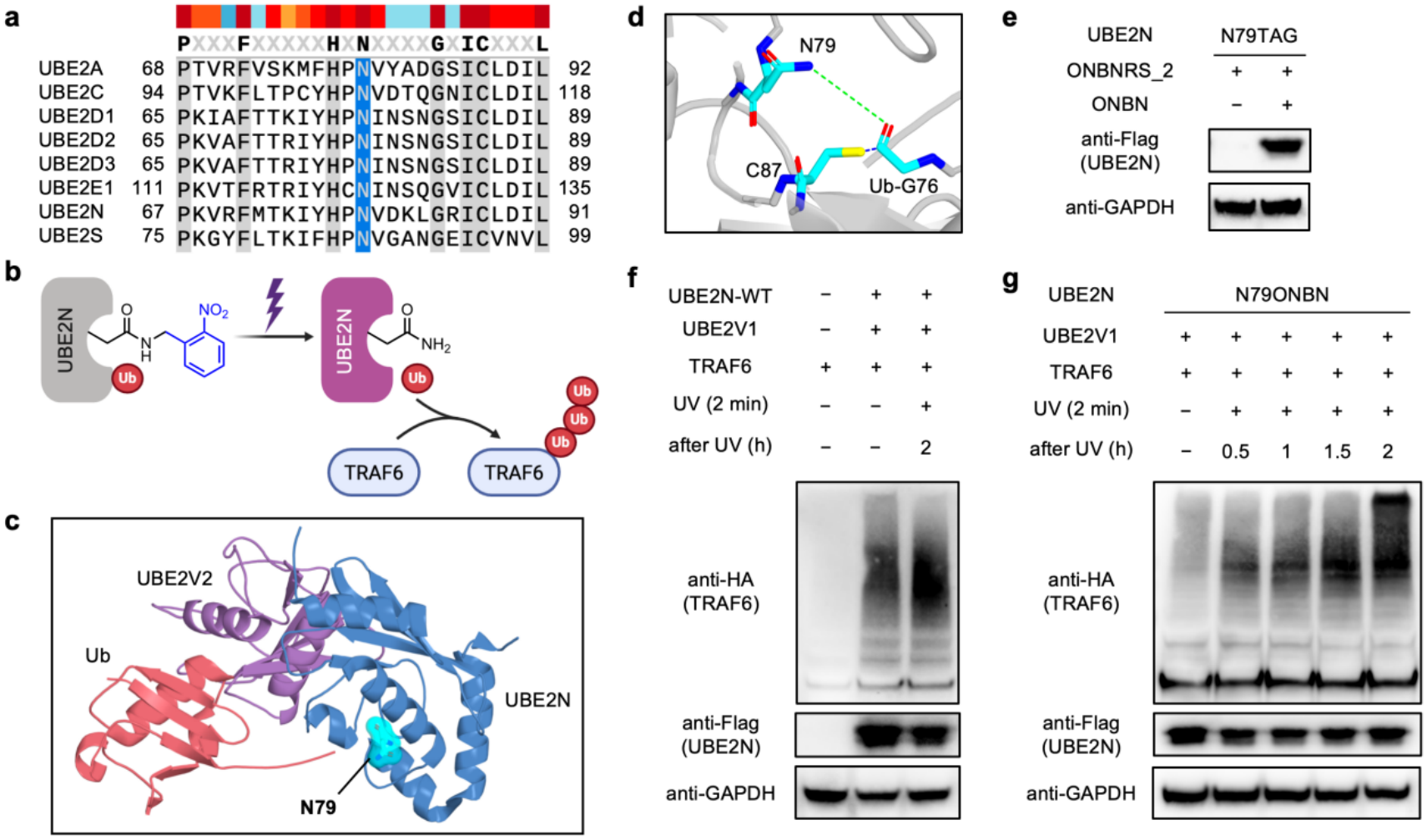
Optical control of UBE2N-mediated ubiquitination through ONBN decaging. **(a)** Human E2 sequence alignment highlighting the conserved Asn in the HPN motif. **(b)** ONBN caging and UV-triggered restoration of UBE2N function in TRAF6 ubiquitination. **(c)** Yeast E2– cofactor–Ub complex (PDB 2GMI)^47^, labelled using UBE nomenclature: UBE2N, blue; UBE2V2 (Mms2), purple; Ub, red. N79 is highlighted by a cyan semitransparent molecular surface. **(d)** Human WT UBE2N from PDB 1J7D^48^ superposed on the yeast E2 chain of 2GMI, retaining the original Ub coordinates. N79, C87 and Ub-G76 are shown as sticks. Green and blue dashed guides connect N79 to the Ub-G76 carbonyl O, and C87 Sγ to the Ub-G76 carbonyl C, respectively. Dashed lines are visual guides in the rigid superposition. **(e)** ONBN-dependent UBE2N-N79TAG expression with ONBNRS_2 in HEK293T cells. **(f)** TRAF6 immunoblots with or without WT UBE2N and UBE2V1, including a sample collected 2 h after 2 min UV irradiation. **(g)** TRAF6 immunoblots in cells expressing UBE2N-N79ONBN and UBE2V1, collected at 0.5, 1, 1.5 and 2 h after 2 min UV irradiation, alongside a nonirradiated control. Anti-HA detects TRAF6, anti-Flag detects UBE2N and GAPDH is the loading control. Immunoblots are representative of three independent experiments.

To examine this strategy in cells, HEK293T cells were cotransfected with ONBNRS_2/tRNA and a P2A-linked construct encoding Flag-tagged UBE2N-N79TAG, UBE2V1 (Uev1a, a noncatalytic cofactor for UBE2N) and HA-tagged TRAF6. Full-length UBE2N was detected with ONBN supplementation, confirming ONBN-dependent amber suppression (Fig. 3e; Fig. S12). TRAF6 polyubiquitination in the WT control showed no apparent change after UV irradiation (Fig. 3f). We next examined the cellular ubiquitination response following optical activation. Cells expressing UBE2N-N79ONBN were irradiated for 2 min and harvested after 0.5, 1, 1.5 and 2 h. Compared with the nonirradiated control, irradiated samples showed stronger high-molecular-weight TRAF6-reactive ubiquitination bands (Fig. 3g; WT/N79A controls and cofactor comparisons, Figs. S12, S13). These results support ONBN-mediated photocaging of N79 as a strategy for optical control of UBE2N-mediated TRAF6 polyubiquitination in living cells. By providing a defined optical trigger, this approach enables monitoring of subsequent cellular ubiquitination responses and may facilitate future investigations of ubiquitination dynamics and UBE2N-dependent substrate modification.

### Development of CAGE-prox^N^ for optical control of firefly luciferase

Having established that photocaging of functionally important Asn residues enables direct regulation of protein activity, we next explored whether ONBN could control enzyme activity when introduced at noncatalytic positions near the active site. The CAGE-prox concept has also been extended to on-demand protein activation in living mice^49^. Inspired by the CAGE-prox strategy^20^, we reasoned that the bulky photoremovable group of ONBN could interfere with substrate accommodation through local steric effects. Photodecaging would remove this group, leaving an uncaged Asn residue at the engineered position and potentially relieving the inhibition. This design would extend Asn-based photocontrol beyond the direct masking of native functional Asn residues. We therefore investigated an Asn-based implementation of proximal decaging, termed CAGE-prox^N^.

We selected firefly luciferase (FLuc), which catalyzes the ATP-dependent oxidation of D-luciferin^50^, as a model enzyme to evaluate CAGE-prox^N^, with ONBN-containing variants expressed using ONBNRS_1 (Fig. 4a). We first modeled the FLuc–luciferyl-AMP complex using AlphaFold 3 and refined the predicted structure with Rosetta Relax for subsequent site selection (Fig. 4b; pre-relaxation model, Fig. S14). Candidate positions near the substrate-binding pocket were identified using expanded CAGE-prox geometric criteria^20^ describing their distance and orientation relative to the ligand (distance below 6.5 Å and angle below 120°). Screening all 550 residues retained 59 positions (Fig. 4c). We then computationally evaluated substitution of the original residue with uncaged Asn using Rosetta^51^, assessing predicted changes in protein stability to prioritize positions compatible with the post-decaging state. Folding-score changes for the 59 geometrically selected positions were plotted against FLuc residue number; a cutoff of Δ*G*_*f*_ < 0.2 REU retained ten positions (Fig. 4d). We experimentally assayed all ten folding-filter positions as ONBN-incorporation variants, together with WT FLuc. Lysates were irradiated for 2 min, and comparison of the +UV signal with the +UV/−UV activation ratio identified A348 as the strongest photoresponsive candidate, with a 29.75-fold increase in mean luminescence (Fig. 4e; Fig. S15). At A348, luminescence increased with UV exposure of intact cells over 0–180 s, followed by immediate harvest and a lysate assay (Fig. 4f; Fig. S16; *n* = 4 per time point). Live-cell luminescence imaging and immunoblotting provide the corresponding exposure series (Fig. 4g,h). These results demonstrate light-dependent activation of FLuc through ONBN incorporation at an engineered proximal position, supporting the feasibility of extending Asn-based photocaging to proximal decaging.

**Figure 4.**
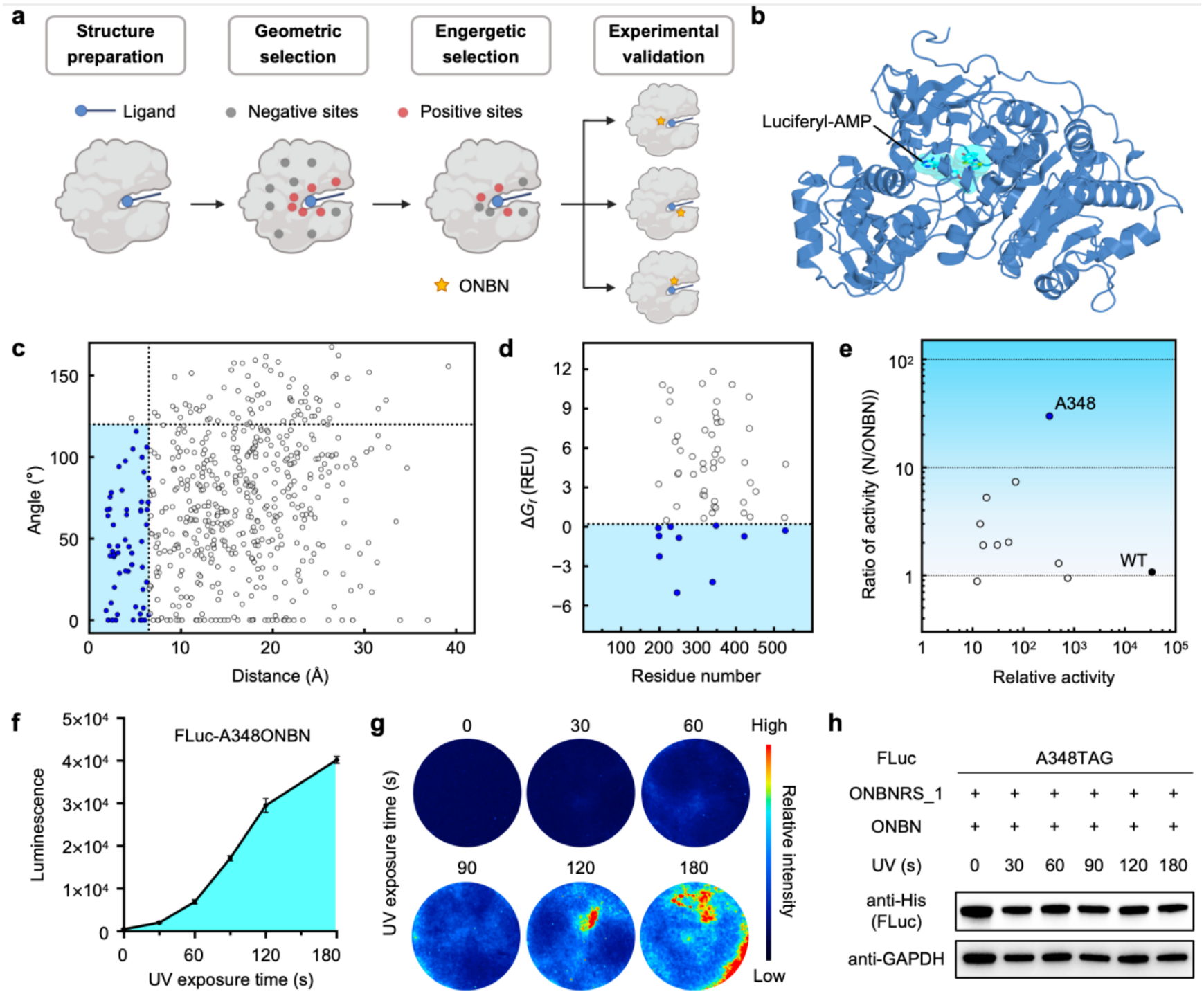
CAGE-prox^N^-guided optical control of firefly luciferase. **(a)** Structure preparation, geometric and energetic selection, and experimental validation of CAGE-prox^N^. **(b)** Final relaxed FLuc–luciferyl-AMP model. **(c)** Geometric screening of 550 residues; 59 satisfy a distance below 6.5 Å and an angle below 120° (geometric selection as described in the text). **(d)** Rosetta folding-score changes (Δ*G*_*f*_) for the 59 geometrically selected positions versus FLuc residue number, shown on linear axes. The dotted line marks Δ*G*_*f*_ = 0.2 REU; ten positions below the cutoff are blue^51^. **(e)** Ten ONBN-incorporation sites and WT FLuc: *x* = mean(+UV signal)/10 and *y* = mean(+UV)/mean(−UV), with three replicates per condition and logarithmic axes. A348 is blue; WT is black. Lysates were irradiated for 2 min. **(f)** FLuc-A348ONBN activity after UV irradiation of intact cells and immediate harvest (mean ± SEM, *n* = 4). **(g)** Live-cell luminescence at the same exposure times. **(h)** Anti-His immunoblots of FLuc-A348ONBN after the indicated exposures and immediate harvest; GAPDH is the loading control. Immunoblots are representative of three independent experiments.

### Optical control and kinetic analysis of post-translational N-glycosylation

Having demonstrated ONBN-mediated optical control of enzyme function at both functional Asn residues and noncatalytic positions near active sites, we next turned to another important biological role of Asn: serving as the acceptor residue for N-linked glycosylation. This modification contributes to the folding, stability and function of secretory and membrane proteins^31,52,53^. In mammalian cells, glycan transfer occurs predominantly during protein synthesis and translocation into the endoplasmic reticulum (ER), whereas a subset of acceptor sites undergoes post-translational modification, often mediated by the STT3B-containing oligosaccharyltransferase complex (OST-B)^54,55^. The coupling of glycosylation to protein biogenesis complicates the analysis of how local sequence context influences the time course of glycan installation. To temporally separate glycosylation initiation from reporter expression and maturation, we constructed ER-targeted EGFP reporters bearing an N-glycosylation sequon within a flexible C-terminal extension and incorporated ONBN at the acceptor Asn position (Fig. S17). Photodecaging restores the native Asn side chain, allowing the glycan-acceptor site to be activated at a defined time in living cells. Using this platform, we investigated both the temporal progression of glycosylation at a single site and the influence of different local sequence contexts on glycosylation kinetics (Fig. 5a).

**Figure 5.**
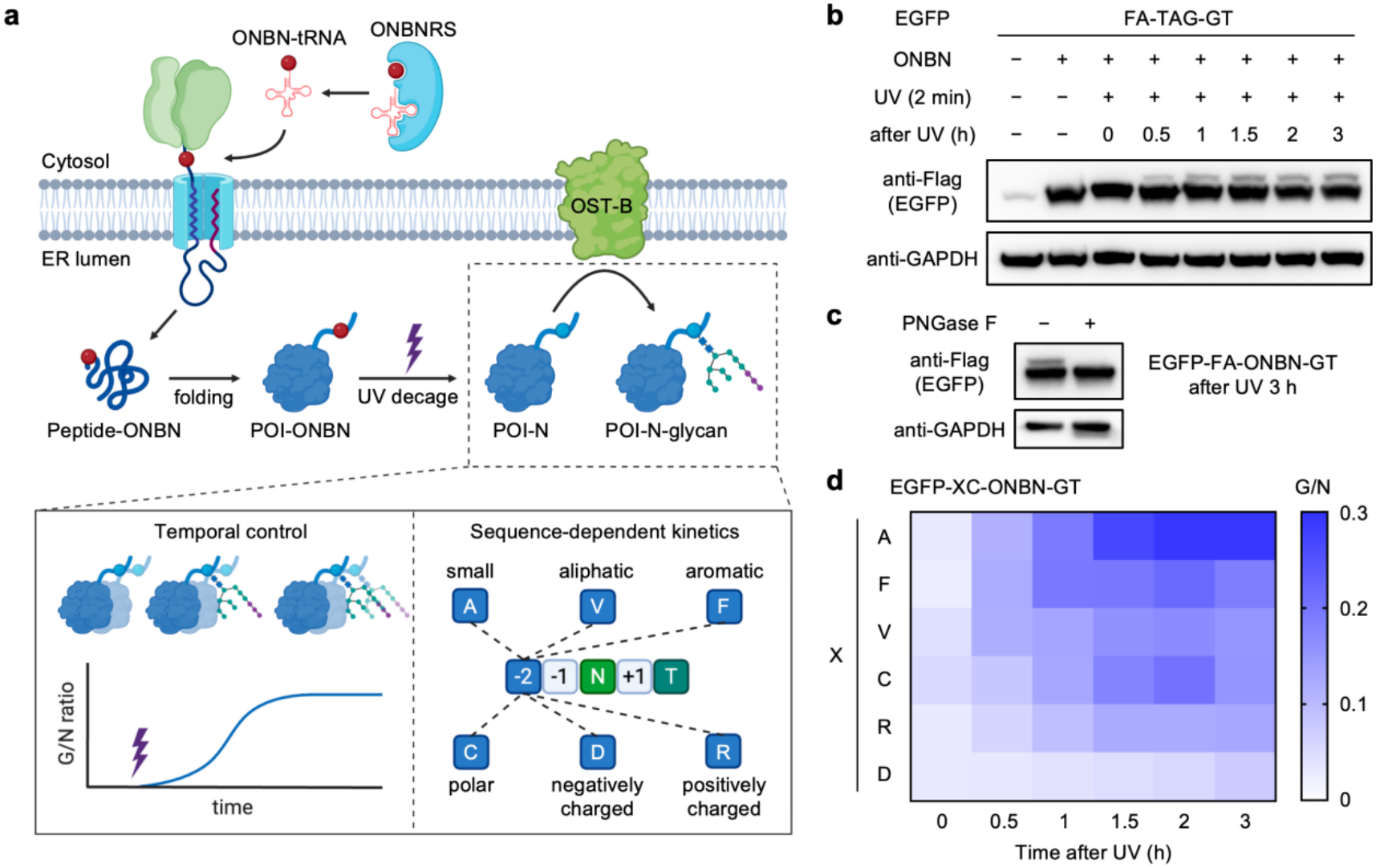
Optical control of N-glycosylation in living cells. **(a)** ONBN incorporation into an ER-targeted EGFP reporter and light-triggered activation of its glycan-acceptor Asn. OST-B is depicted schematically as a proposed mediator. Time-resolved sampling and variation of the −2 residue probe glycosylation kinetics. **(b)** Anti-Flag immunoblots of the FANGT-derived EGFP-FA-ONBN-GT reporter at 0, 0.5, 1, 1.5, 2 and 3 h after 2 min UV irradiation, with the indicated nonirradiated and ONBN-omission controls. **(c)** PNGase F treatment of the FANGT-derived reporter collected 3 h after irradiation. GAPDH is the loading control in b and c. **(d)** Heatmap of glycosylated-to-nonglycosylated EGFP intensity ratios (G/N) for the XCNGT series. Rows show X = A, F, V, C, R and D at the −2 position; columns show 0, 0.5, 1, 1.5, 2 and 3 h. The −1 residue is Cys in this series and Ala in the FANGT reporter. ONBN initially replaces the acceptor Asn. Each cell is the mean of three G/N measurements on a common 0–0.3 scale, without row-wise normalization. Immunoblots and complete time courses for all nine XCNGT variants are in Figure S19. Immunoblots are representative of three independent experiments.

We constructed glycosylation reporters containing FANGT or WCNGT, two previously reported enhanced aromatic sequons^32,33^ (Fig. S18). We first characterized the glycosylation time course of the FANGT reporter, with ONBN replacing the acceptor Asn through genetic code expansion. HEK293T cells were cultured with 1 mM ONBN for 48 h, after which the medium was replaced with ONBN-free medium before UV irradiation for 2 min. Photodecaging regenerated the native Asn-containing motif, making it available for recognition by cellular OST. Samples collected at successive time points after the end of irradiation were analyzed by immunoblotting to monitor the appearance of the glycosylated reporter (Fig. 5b). A higher-molecular-weight band became evident after photodecaging, and its relative abundance increased over the 3 h time course. To determine whether this mobility shift resulted from N-glycosylation, samples collected 3 h after irradiation were treated with PNGase F. Treatment eliminated the upper band and collapsed the reporter signal to the lower-molecular-weight species, confirming that the observed shift reflected N-linked glycan attachment (Fig. 5c). These results establish optical control over glycosylation initiation following reporter expression and enable time-resolved monitoring of glycan installation in living mammalian cells.

We next examined how residues flanking the acceptor sequon influence glycosylation kinetics. Previous studies have shown that the residue two positions N-terminal to the acceptor Asn (the −2 position) can modulate glycosylation efficiency^30,32,33^. We therefore varied this position within a common XCNGT reporter scaffold, retaining Cys at the −1 position and the NGT core sequon, generating a panel of contexts with distinct side-chain properties. Glycosylation was monitored after photodecaging, and glycosylated-to-nonglycosylated EGFP ratios (G/N) were calculated by immunoblot densitometry to compare the time-dependent accumulation of glycosylated reporter (Fig. 5d; Fig. S19). Reporters bearing Ala or Phe at the −2 position exhibited the most rapid increases in G/N, whereas those containing the charged residues Asp or Arg showed the slowest increases among the variants examined. These findings demonstrate that residues outside the canonical N-glycosylation sequon influence the apparent kinetics of glycan installation in cells. By combining a defined activation time with a common substrate scaffold, ONBN photodecaging enables local sequence effects to be resolved at the level of glycosylation time courses, providing a means to investigate the kinetic determinants of cellular N-glycosylation.

## Conclusions

We have established a genetically encoded photocaging strategy that enables light-triggered regeneration of the asparagine side-chain carboxamide in proteins. To the best of our knowledge, this is the first genetically encoded decaging strategy targeting a native carboxamide side chain. ONBN incorporation through engineered synthetase/tRNA pairs supports optical activation of *Renilla* luciferase and UBE2N-mediated TRAF6 polyubiquitination in living cells. CAGE-prox^N^ further extends this approach to an engineered position near the firefly luciferase active site, where photodecaging leaves an Asn residue compatible with enzyme activity. These results establish complementary designs for controlling protein function through native functional Asn residues and introduced proximal residues. Beyond protein activation, ONBN allows the glycan-acceptor function of Asn to be switched on at a defined time. Photodecaging of an acceptor sequon in an ER-targeted reporter separates site activation from reporter expression and maturation, enabling time-resolved analysis of post-translational glycan installation. Differences among reporters bearing distinct −2 residues demonstrate that local sequence context outside the canonical sequon influences apparent glycosylation kinetics in cells. By providing a common optical starting point for these measurements, ONBN offers a means to compare glycosylation time courses across sequence contexts. Together, these capabilities expand the functional-group scope of genetically encoded gain-of-function control and provide a platform for investigating Asn-dependent protein functions and the kinetic determinants of cellular N-glycosylation.

