## Supporting Information for "Genetically Encoded Photodecaging of Asparagine Enables Protein Function Control and Time-Resolved N-Glycosylation"

* Correspondence to: Peng R. Chen

#### Contents


General Information 3

Supplementary Methods 5

1. Chemical synthesis and characterization of ONBN 5
2. Small-molecule photodecaging and UPLC–MS analysis 5
3. Plasmid construction and sequence verification 6
4. Prediction of the BnDRS–ONBN–AMP complex and hotspot identification 6
5. Construction and quality control of the NNK library 7
6. Positive and negative selection of ONBNRS variants 7
7. Bacterial reporter expression and fluorescence measurements 7
8. Bacterial sfGFP expression purification and in vitro photodecaging 8
9. Intact-protein mass spectrometry and site-specific incorporation analysis 8
10. Mammalian cell culture and transfection 9
11. UV irradiation and ONBN cellular compatibility 9
12. General immunoblotting and source-image preparation 10
13. EGFP fluorescence measurements and immunoblot validation of incorporation 10
14. Mammalian EGFP expression enrichment and live-cell photodecaging 11
15. Optical activation and luminescence measurement of *Renilla* luciferase 11
16. Sequence alignment and structural analysis of E2 enzymes 12
17. Optical activation of UBE2N and analysis of TRAF6 ubiquitination 12
18. FLuc complex modeling and CAGE-prox$^N$ site selection 13
19. Experimental evaluation of proximal FLuc sites and UV dose response 13
20. ER reporter design and time-resolved glycosylation 14
21. PNGase F validation of N-linked glycosylation 15
22. Sequon-dependent kinetics and immunoblot densitometry 15

Supplementary References 16

Supplementary Figures S1–S3 and S5–S19 17

NMR Spectra 36

High Resolution Mass Spectra 38


The exact positions and residue identities of the synthetase variants are omitted from this version.

### General Information

#### Materials

ONBN was synthesized as described in Supplementary Method 1. Organic synthesis reagents were obtained from Xinhengyan, BLD Pharmatech and Beijing Tongguang Fine Chemical: Boc-Asp-OtBu (BLD Pharmatech, CAS 34582-32-6), 2-nitrobenzylamine hydrochloride (Xinhengyan), EDCI (BLD Pharmatech, CAS 25952-53-8), HOBt (BLD Pharmatech), triethylamine (Beijing Tongguang Fine Chemical) and 4 M HCl in 1,4-dioxane (Beijing Tongguang Fine Chemical). Solvents (ethyl acetate, petroleum ether, 1,4-dioxane) were from Beijing Tongguang Fine Chemical and Concord Technology. Column chromatography used silica gel (Yinlong, A-200300-P, 200–300 mesh) on a Cheetah Pro system (Agela) with SepaFlash prepacked columns (Santai Technologies).

DNA amplification used 2× Phanta Flash Master Mix (Vazyme, P510). Library assembly used BsaI (Beyotime, D6128) and T4 DNA ligase (NEB, M0202). Seamless assembly used ClonExpress Ultra One Step Cloning Kit V3 (Vazyme, C117). PCR products were purified with a SanPrep Column PCR Product Purification Kit (Sangon Biotech, B518141-0100). Plasmid DNA was isolated with an Omega Bio-tek Endo-free Plasmid Mini Kit II (D6950-01). Ampicillin (Solarbio, A8180), chloramphenicol (Solarbio, C8050) and kanamycin (Solarbio, K8020) were used at final concentrations of 100, 37 and 50 μg $mL^{-1}$, respectively. Each antibiotic stock was diluted 1:1,000: ampicillin, 100 mg $mL^{-1}$ in water; chloramphenicol, 37 mg $mL^{-1}$ in ethanol; and kanamycin, 50 mg $mL^{-1}$ in water. These concentrations applied to plates, colony starter cultures, plasmid amplification and induced expression unless otherwise stated. L-Arabinose (Xinhengyan, Z005524-500g) was used for induction; agar (Xinhengyan) was used for plates and agarose (Yeasen, 10208ES60) for gel electrophoresis. LB medium was prepared with yeast extract (Oxoid, LP0021B) and tryptone (Oxoid, LP0042).

Lipo8000 (Beyotime, C0533) was used for transfection, routinely in 12-well plates. Samples for immunoblotting were lysed in Western and IP lysis buffer (Beyotime, P0013) supplemented with a 1× protease inhibitor cocktail freshly diluted 1:100 from the Beyotime 100× stock (P1005). RLuc and FLuc activity samples were lysed using the buffers supplied with their respective assay kits. Myc-tagged EGFP was enriched with Myc-Tag IP magnetic beads and eluted with the supplied Myc peptide (Beyotime, P2183M). Blocking used QuickBlock (Beyotime, P0252-500ml). PBS (Solarbio, P1020) and 10× TBST (Solarbio, T1081) were used for washing and buffer preparation. His-tagged proteins were purified on 5 mL nickel-affinity columns (Yeasen, 20504ES) using an ÄKTA start system; buffer exchange and size-exclusion chromatography used Amicon Ultra centrifugal filters (Millipore; UFC900396, 15 mL, 3 kDa; UFC800396, 4 mL, 3 kDa; UFC5003BK, 0.5 mL, 3 kDa) and an ÄKTA pure 25 system (Cytiva) equipped with a Super75 column (NanoMicro; 11 × 300 mm) and controlled with UNICORN 7.0.

Immunoblotting used SurePAGE 4–20% precast gels (GenScript; M00655, 10-well; M00657, 15-well) and 20× MOPS running buffer (GenScript, M00680-500). Glycosylation reporters were resolved on 12% SurePAGE gels (GenScript, M00669). Proteins were transferred with NCM Biotech rapid transfer buffer (WB4600) onto 0.2 μm PVDF membranes (Millipore, Immobilon-PSQ, ISEQ00010). For nonreducing UBE2N–Ub samples, 4× NuPAGE LDS Sample Buffer (Thermo Fisher Scientific, NP0007) was used at 1× without reducing reagent. Reducing denaturing loading buffer (CWBIO, CW0027S) and stripping buffer (CWBIO, CW0056M) were used as supplied. Molecular-weight standards were the 180 kDa Plus Prestained Protein Marker (Vazyme, MP201-01) and PageRuler Prestained Protein Ladder (Thermo Fisher Scientific, 26616).

*Renilla* luciferase activity was measured with the *Renilla* Luciferase Reporter Gene Assay Kit (Beyotime, RG016). Firefly luciferase activity was measured with the Firefly Luciferase Reporter Gene Assay Kit (Yeasen, 11401). MTT absorbance was measured with the MTT Cell Proliferation and Cytotoxicity Assay Kit (Elabscience, E-CK-A341) with absorbance read at 570 nm. PNGase F treatment used the PNGase F Deglycosylation Kit (Beyotime, P2318S). D-Luciferin sodium salt (Yeasen, 40901ES01) was diluted from a 100 mM stock to 1 mM in PBS for live-cell FLuc imaging. Where indicated, MG132 (Beyotime, S1748-5mg) was used at the stated concentration.

The cell line used for mammalian experiments was HEK293T. Cells were maintained in high-glucose DMEM (Gibco, 11965092) supplemented with 10% fetal bovine serum (VivaCell, C04001-500) and 1× penicillin–streptomycin solution (Vivacell, C3420-0100) at 37 °C in 5% $CO_2$. Trypsin-EDTA (0.25%, Vivacell, C3530-0100) was used for passaging, and Opti-MEM (Gibco, 31985070) for transfection. *E. coli* DH5α was used for routine cloning, library-plasmid amplification and plasmid preparation. *E. coli* DH10B was used for induced reporter expression and ONBNRS screening. Primary antibodies were CST monoclonal antibodies: anti-Myc (9B11, mouse, #2276), anti-His (D3I1O, rabbit, #12698), anti-Flag (9A3, mouse, #8146), anti-HA (C29F4, rabbit, #3724) and anti-GAPDH (14C10, rabbit, #2118). HRP-linked anti-mouse (#7076) and anti-rabbit (#7074) secondary antibodies were from CST. Antibodies were used at 1:1,000 (primary) and 1:10,000 (secondary) unless stated otherwise.

**Instruments**

$^1H$ and $^{13}C$ NMR spectra were recorded on a Bruker Avance spectrometer at 500 and 126 MHz, respectively. High-resolution mass spectra were recorded on a Bruker Solarix XR FTMS mass spectrometer using positive-mode electrospray ionization (ESI) with methanol as the solvent. Data were analyzed using Bruker Compass DataAnalysis 5.0. Chemical shifts are reported in ppm relative to the residual solvent signal. Multiplicities are designated s, singlet; d, doublet; t, triplet; q, quartet; dd, doublet of doublets; and m, multiplet.

UPLC–MS analyses used an ACQUITY UPLC system with a photodiode-array detector and a Waters SQ Detector 2 (Waters). The ONBN photolysis series in Figure S1 used a BEH C18 column (1.7 µm; Waters) at 40 °C with a 5 µL injection. Intact-protein analyses used 80 °C and a 5 µL injection. Absorbance, fluorescence and luminescence were measured on a Synergy H1 multi-mode microplate reader (BioTek) with Gen5 3.11.19 software. Luminescence measurements used top optics, unfiltered emission, a 1 s integration time, a 100 ms delay, extended dynamic range and a read height of 4.5 mm. Gain was 200 for the final FLuc site screen, the detailed FLuc-A348 time course and the expression-adjusted RLuc assay in Figure 2f, and 180 for the unadjusted RLuc dataset in Figure S11a. Bacterial sfGFP fluorescence used 488 nm excitation, 550 nm emission, gain 100, a 7 mm read height, a 100 ms delay and 10 measurements per data point. Cellular EGFP fluorescence used top optics, gain 100, 488 nm excitation, 550 nm emission, a 7 mm read height, a 100 ms delay and 10 measurements per data point. Nucleic acid and protein concentrations were measured on a NanoDrop 2000 spectrophotometer (Thermo Fisher Scientific). Fluorescence microscopy used a Leica DMi8 inverted microscope equipped with an N PLAN 10×/0.25 dry objective, a GFP filter cube and a DFC420C camera, controlled with LAS X 3.3.0.16799. Gel and immunoblot images were acquired on a ChemiDoc XRS+ imaging system (Bio-Rad). Photodecaging used a UVGO 365 nm curing lamp (80 W) at an irradiance of 350 mW $cm^{-2}$ at the sample plane.

PCR was performed on a RePure-T thermal cycler (BIO-GENER). Flash chromatography used a Cheetah Pro system (Agela Technologies) with SepaFlash prepacked columns (Santai Technologies). Solvents were removed on a Heidolph rotary evaporator connected to a WELCH VCZ 524 diaphragm pump and a DL-400 coolant circulation pump (Zhengzhou Saimeide). Nickel-affinity purification used an ÄKTA start system, and desalting and size-exclusion chromatography used an ÄKTA pure 25 system (Cytiva) equipped with a Super75 column (NanoMicro; 11 × 300 mm) and controlled with UNICORN 7.0. TLC used silica gel plates with petroleum ether/ethyl acetate (1:1, v/v) as the mobile phase.

Antibodies and affinity reagents, including suppliers and working dilutions, are listed in Table S1D of the separate Supplementary Data document.

#### Supplementary Methods

##### 1. Chemical synthesis and characterization of ONBN

$N^4$-(2-Nitrobenzyl)-L-asparagine (ONBN) was synthesized by coupling the side-chain carboxyl group of $N^{\alpha}$-Boc-L-aspartic acid α-*tert*-butyl ester (Boc-Asp-OtBu) with 2-nitrobenzylamine, followed by removal of both protecting groups to afford ONBN hydrochloride.

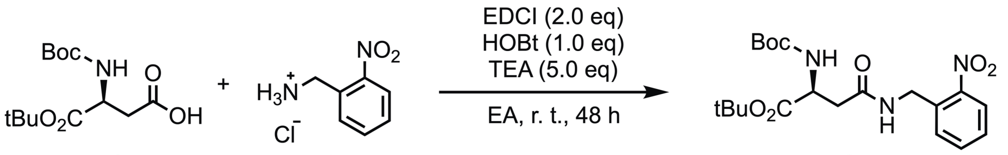


1 2 (1.5 eq) 3 76.5%

A mixture of Boc-Asp-OtBu **1** (15 mmol, 1.0 equiv), 2-nitrobenzylamine hydrochloride **2** (22.5 mmol, 1.5 equiv), EDCI (30 mmol, 2.0 equiv), HOBt (15 mmol, 1.0 equiv) and triethylamine (75 mmol, 5.0 equiv) in ethyl acetate (250 mL) was stirred at room temperature for 48 h. The mixture was concentrated under reduced pressure and dry-adsorbed onto silica gel. Purification was performed on a Cheetah Pro automated flash chromatography system (Agela) with a SepaFlash prepacked column (Santai Technologies), using a gradient of ethyl acetate in petroleum ether to 40% ethyl acetate, followed by an isocratic hold at petroleum ether/ethyl acetate 3:2 (v/v). Fractions were monitored by TLC (petroleum ether/ethyl acetate, 1:1, v/v; $R_f$ = 0.3–0.4). Product-containing fractions were combined, concentrated by rotary evaporation and dried under vacuum to give $N^{\alpha}$-Boc-ONBN *tert*-butyl ester **3** (yield: 76.5%). $^1$H NMR (500 MHz, $CDCl_3$) δ 8.06 (dd, $J$ = 8.3, 1.3 Hz, 1H), 7.69 – 7.55 (m, 2H), 7.46 (ddd, $J$ = 8.7, 7.1, 1.8 Hz, 1H), 6.48 (s, 1H), 5.66 (d, $J$ = 8.3 Hz, 1H), 4.67 (d, $J$ = 6.4 Hz, 2H), 4.48 – 4.26 (m, 1H), 2.91 – 2.64 (m, 2H), 1.41 (d, $J$ = 12.6 Hz, 18H). $^{13}$C NMR (126 MHz, $CDCl_3$) δ 170.24, 155.69, 148.34, 134.08, 133.49, 132.32, 128.77, 125.10, 82.16, 79.79, 50.98, 41.17, 38.30, 28.31, 27.85. HRMS (ESI+) *m/z*: calcd for $C_{20}H_{30}N_3O_7$ $[M+H]^+$ 424.2078; found 424.2083.

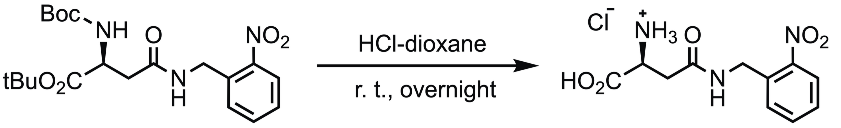


3 4 94.4%

Compound **3** was dissolved in 4 M hydrogen chloride in 1,4-dioxane (25 mL per gram of intermediate) and stirred overnight at room temperature. The resulting white precipitate was collected by centrifugation at 4,000 × g for 30 min and washed three times with ethyl acetate. The precipitate was transferred to a round-bottom flask using methanol, concentrated by rotary evaporation and dried under vacuum to afford ONBN hydrochloride **4** as a white solid (yield: 94.4%). $^1$H NMR (500 MHz, $D_2O$) δ 7.97 (dd, $J$ = 8.2, 1.3 Hz, 1H), 7.57 (td, $J$ = 7.6, 1.3 Hz, 1H), 7.44 – 7.36 (m, 2H), 4.56 (d, $J$ = 2.2 Hz, 2H), 4.11 (td, $J$ = 5.6, 1.8 Hz, 1H), 2.87 (d, $J$ = 5.6 Hz, 2H). $^{13}$C NMR (126 MHz, $D_2O$) δ 171.11, 170.96, 147.57, 134.36, 132.54, 130.07, 128.79, 125.23, 49.72, 40.99, 34.32. HRMS (ESI+) *m/z*: calcd for $C_{11}H_{14}N_3O_5$ $[M+H]^+$ 268.0928; found 268.0936.

##### 2. Small-molecule photodecaging and UPLC–MS analysis

ONBN was dissolved in PBS (pH 7.4) at 1 mM, and 50 µL aliquots were placed in the wells of a 96-well plate. For Figure S1, samples were irradiated at 365 nm for 0, 30 or 60 s. Dark controls were processed in parallel. Samples were protected from further light exposure immediately after irradiation and analyzed by UPLC–MS.

UPLC–MS was performed on an ACQUITY UPLC system equipped with a photodiode-array detector and a Waters SQ Detector 2 (Waters), using a BEH C18 column (1.7 µm; Waters) at 40 °C. The injection volume for this series was 5 µL. Solvent A was acetonitrile (Fisher Scientific, A955-4), and solvent B was water/acetonitrile (95:5, v/v). At a flow rate of 0.3 mL $min^{-1}$, the gradient used 5% A at 0–0.1 min, a linear increase to 100% A at

3.5 min, a hold at 100% A until 5.0 min, a return to 5% A at 5.1 min and equilibration until 6.0 min. The photodiode-array detector acquired spectra from 190 to 300 nm. Positive- and negative-ion electrospray spectra were acquired over *m/z* 100–800 in continuum mode with a 1 s scan time and a cone voltage of 35 V.

PDA, positive-ion and negative-ion chromatograms are shown in Figure S1a–c. The negative-ion mass spectra in Figure S1d–f were combined over scans 35–51 for the dark sample, 26–40 after 30 s irradiation and 26–41 after 60 s irradiation, as recorded in the native MassLynx exports. The chromatographic changes and associated mass signals were used to assess photolysis.

#### 3. Plasmid construction and sequence verification

Reporter and protein-expression plasmids were generated by PCR-based cloning or site-directed mutagenesis using Phanta Flash Master Mix (Vazyme, P510). A standard 50 µL PCR contained 25 µL 2× master mix, 2 µL of each primer (10 µM), 1 µL template DNA (50–100 ng $\mu L^{-1}$) and 20 µL water. PCR used 95 °C for 3 min; 35 cycles of 95 °C for 10 s, 65 °C for 5 s and 72 °C for 40 s; 72 °C for 1 min; and a 4 °C hold. The heated lid was set to 105 °C. Parental template was removed by adding 1 µL DpnI (Beyotime, D6257S) and 5.6 µL 10× buffer Y, followed by incubation according to the supplier's program. PCR products were purified with a SanPrep Column PCR Product Purification Kit (Sangon Biotech, B518141-0100) before assembly; site-directed mutagenesis products were used directly. For seamless assembly, vector and insert were mixed at 1:1 in a 10 µL reaction containing 5 µL 2× CE Mix V3 (Vazyme, C117) and incubated at 50 °C for 30 min. Assembly products (5 µL) were transformed into chemically competent *E. coli* DH5α: ice for 30 min, heat shock at 42 °C for 60 s, ice for 1 min, recovery in 1 mL LB at 37 °C for 1 h, and plating on LB agar with the appropriate antibiotic. Amber codons were introduced at sfGFP-D190, EGFP-Y40, RLuc-N53, UBE2N (Ubc13) at N79, the candidate FLuc positions and the glycan-acceptor Asn of the ER reporter. Wild-type and uncaged controls were constructed in the same vector background. Constructs were verified by Sanger sequencing (GENEWIZ). Plasmid DNA was prepared with the Omega kit described above.

The bacterial synthetase/tRNA expression vector pSUPAR (pBAD-ONBNRS) carries an araBAD-driven 419-residue ONBNRS, a proK-driven suppressor-tRNA cassette, a chloramphenicol-resistance marker and a p15A origin. The supplied pSUPAR construct encodes ONBNRS_2. Mammalian expression used pRS (pCMV-NES-ONBNRS-4x7sk), with a CMV-driven NES–synthetase fusion, four 7SK-driven tRNA cassettes and a bacterial ampicillin-resistance marker. ONBNRS_2 was used for bacterial sfGFP purification and MS and for the E2 experiments. ONBNRS_1 was used for cellular EGFP photodecaging MS, RLuc, CAGE-prox$^{N}$/FLuc and glycosylation. Both variants were tested in bacterial sfGFP and mammalian EGFP expression assays. The supplied pCMV-NES-ONBNRS-4x7sk DNA record encodes the ONBNRS_1 variant; its annotated vector organization is shown in Figure S3c.

The empty vector pCMV-Vector-A-4x7sk contains four suppressor-tRNA cassettes under 7SK promoters and was the empty-vector counterpart for most mammalian constructs. pCMV-Vector-B was the empty-vector counterpart for FLuc and RLuc. Both backbones contain CMV expression elements and bacterial ampicillin resistance.

#### 4. Prediction of the BnDRS–ONBN–AMP complex and hotspot identification

BnDRS was used as the parental synthetase[1,2] on the 419-residue chPylRS framework[3], with the IPYE substitutions[4]. The BnDRS–ONBN–AMP complex was predicted using the AlphaFold 3 workflow described for PABD[5,6], with BnDRS and ONBN–AMP replacing the corresponding PABD-system components. The saved docking session contains the 419-residue BnDRS protein (chain A), a 72-nucleotide suppressor tRNA (chain B) and ONBN–AMP (chain C). The suppressor tRNA sequence recovered from the session is provided below.

Suppressor tRNA sequence from the saved structural session (5′→3′):

```
UGGCGGAAACCCCGGAAAUCUAACCCGGCUGAACGGAUUUAGAGUCCAUUCGAUCUACAUGAUCAGGUUUCC
```

The predicted ONBN–AMP pocket guided the choice of five positions, defining the focused NNK libraries. Figure S2 shows the overall complex and ligand conformation from the saved session; all views retain the saved atomic coordinates.

#### 5. Construction and quality control of the NNK library

Focused NNK libraries targeting the five positions were constructed in a kanamycin-resistant pBK backbone expressing BnDRS from the oxb20 promoter (Fig. S3a). NNK codons (N = A, T, G or C; K = G or T) were introduced at the specified positions using Phanta Flash Master Mix (Vazyme). Template-matching sequences flanked the randomized codons; BsaI recognition sites and compatible four-base overhangs enabled assembly of the PCR fragments. Distant randomized positions were introduced on separate fragments. PCR used the 50 µL reaction format and cycling program in Method 3. Amplicons were examined by agarose gel electrophoresis, treated with DpnI and column-purified before assembly.

Parental template was removed from 50 µL pooled PCR product by adding 5.6 µL Buffer Y and 1 µL DpnI, followed by column purification. Fragments were assembled at an insert:vector molar ratio of approximately 5:1. The 40 µL assembly contained 4 µg vector, 2 µg insert, 6 µL BsaI (Beyotime), 4 µL T4 DNA ligase (NEB), 4 µL T4 ligase buffer and water to volume. Restriction–ligation cycling consisted of 110 cycles of 37 °C for 5 s and 16 °C for 4 min 35 s, followed by 60 °C for 10 min, 80 °C for 10 min and a 4 °C hold. For each transformation, 10 µL assembly mixture was added to 100 µL chemically competent *E. coli* DH5α. Heat shock followed Method 3, including recovery in 1 mL LB at 37 °C for 1 h. The recovered cultures were grown with kanamycin for library-plasmid preparation using the Omega kit. Pooled-library Sanger traces were aligned to the synthetase coding sequence to verify randomization at the five positions.

#### 6. Positive and negative selection of ONBNRS variants

*E. coli* DH10B was cotransformed by heat shock with the kanamycin-resistant pBK synthetase library and the ampicillin-resistant positive-selection reporter plasmid A1-pos-Cam-D112TAG-GFPuv-D190TAG-PylT (Fig. S3). Cells recovered in 1 mL LB at 37 °C with shaking for 1 h before plating. Positive-selection LB agar contained ampicillin (100 µg $mL^{-1}$), chloramphenicol (37 µg $mL^{-1}$), kanamycin (50 µg $mL^{-1}$), 0.2% (w/v) L-arabinose and 1 mM ONBN, matching the inducer and ONBN concentrations used for liquid-culture expression. Plates were incubated at 37 °C for 24 h and then at room temperature for a further 48 h before colony picking. Chloramphenicol resistance and GFPuv fluorescence served as amber-suppression readouts. The reporter carries chloramphenicol acetyltransferase (CAT) with D112TAG under a constitutive Ptrp promoter, L-arabinose-inducible GFPuv with D190TAG and a C-terminal $His_6$ tag, and an lpp-driven PylT suppressor-tRNA cassette (Fig. S3b).

Barnase-based negative selection was performed using synthetase plasmids recovered from positive selection. These plasmids were electroporated into *E. coli* DH10B carrying the Barnase_Q2TAG_D44TAG reporter. Cells were recovered in LB at 37 °C for 1 h and plated on LB agar containing 0.1% (w/v) L-arabinose, kanamycin (50 µg $mL^{-1}$) and ampicillin (50 µg $mL^{-1}$), without ONBN. Plates were incubated at 37 °C for 48 h. Colonies were harvested for synthetase-plasmid recovery and reporter validation.

Surviving clones were isolated, retransformed as appropriate, and assessed for reporter expression with and without ONBN. Candidate hits were sequenced and assigned the names ONBNRS_1 and ONBNRS_2 according to their verified amino acid sequences.

#### 7. Bacterial reporter expression and fluorescence measurements

*E. coli* DH10B was cotransformed with the bacterial ONBNRS/tRNA vector and the pBAD-sfGFP-D190TAG reporter. WT chPylRS, BnDRS, ONBNRS_1 and ONBNRS_2 were compared in parallel. Cultures were grown in LB with the appropriate antibiotics; the expression-vector pair carries chloramphenicol and ampicillin resistance, respectively. At $OD_{600}$ = 0.6, L-arabinose and ONBN were added to final concentrations of 0.2% (w/v; 20% stock diluted 1:100) and 1 mM (100 mM stock diluted 1:100), respectively, where indicated. Expression proceeded for 24 h at 25 °C. The Figure 1g groups were −Ara/−ONBN, +Ara/−ONBN and

+Ara/+ONBN. Four culture tubes were grown from the same single-colony clone for each condition. For plate-reader measurements, 200 µL induced bacterial culture was transferred to each well of a conical-bottom 96-well plate and centrifuged at 4,000 × g for 15 min at room temperature. The supernatant was discarded, and fluorescence of the cell pellet was measured on the Synergy H1 reader at 488 nm excitation and 550 nm emission using top optics and a high-energy xenon flash lamp. The reader settings were gain 100, a 7 mm read height, a 100 ms delay and 10 measurements per data point. $OD_{600}$ was also measured. Fluorescence data are summarized as mean ± SD ($n$ = 4).

#### 8. Bacterial sfGFP expression purification and in vitro photodecaging

His-tagged sfGFP-D190ONBN for bacterial protein purification and MS was expressed in *E. coli* DH10B carrying ONBNRS_2 and pBAD-sfGFP-D190TAG. A 20 mL overnight starter culture was expanded into 500 mL antibiotic-supplemented LB and grown at 37 °C to $OD_{600}$ = 0.6. ONBN and L-arabinose were added to 1 mM and 0.2% (w/v), respectively, and expression proceeded for 24 h at 25 °C. Cells were harvested at 4,000 × g for 20 min and resuspended in 20 mL PBS containing 1× protease inhibitor cocktail (Beyotime).

Bacterial suspensions were disrupted by chilled high-pressure homogenization at approximately 1,000 bar. Lysates were clarified at 20,000 × g for 35 min at 4 °C and filtered through a 0.22 µm membrane. The supernatant was loaded onto a 5 mL nickel-affinity column (Yeasen, 20504ES) on an ÄKTA start system. Stepwise washes used NTA0, NTA10 and NTA20, followed by elution with NTA250 and NTA500; NTA denotes the imidazole concentration in mM, and NTA0 is PBS without imidazole. Protein-containing eluates were exchanged into PBS by size-exclusion chromatography on a Super75 column (NanoMicro; 11 mm internal diameter × 300 mm) using an ÄKTA pure 25 system controlled with UNICORN 7.0. Elution was monitored at 280 nm, and fractions with high absorbance were collected separately and selected for centrifugal concentration using Amicon Ultra filters. Samples were then subjected to intact-protein UPLC–MS (Method 9).

For in vitro photodecaging, purified sfGFP-D190ONBN was diluted in PBS to 0.5 mg $mL^{-1}$ and dispensed into a 96-well plate at 50 µL per well. The plate was kept in an ice-water bath and irradiated at 365 nm for 0, 30, 60, 90, 120 or 180 s. Samples were collected after irradiation and submitted directly for UPLC–MS at 80 °C with 5 µL injections, without further purification. Raw charge-state envelopes and MaxEnt1-deconvoluted spectra were retained for each time point. These spectra derive from bacterial sfGFP-D190ONBN; the cellular EGFP-Y40ONBN series was prepared by irradiating intact cells before harvest and Myc enrichment (Method 14).

#### 9. Intact-protein mass spectrometry and site-specific incorporation analysis

Intact-protein mass spectra were acquired on an ACQUITY UPLC system coupled to a Waters SQ Detector 2 (Waters). The column temperature was 80 °C and the injection volume was 5 µL. Mobile phase A was acetonitrile containing 0.1% (v/v) formic acid, and mobile phase B was water containing 0.1% (v/v) formic acid. Raw charge-state envelopes were deconvoluted using MaxEnt1 in MassLynx (Waters). Bacterial sfGFP-D190ONBN was nickel-purified, desalted on ÄKTA pure and concentrated (Method 8); cellular EGFP-Y40ONBN was Myc-enriched and injected directly from peptide eluate without ultrafiltration or solvent exchange (Method 14). Theoretical masses were calculated from the exact encoded sequences, tags, GFP chromophore maturation state and the terminal-processing assumptions specified below.

The cellular EGFP-Y40ONBN construct encodes 249 residues including the C-terminal Myc tag. Average neutral masses were calculated from this exact sequence, replacing residue 40 with Asn for the decaged species. For the N-terminally acetylated, initiator-Met-cleaved proteoform, chromophore maturation subtracts $H_2O$ + $H_2$ (20.031 Da), initiator-Met removal subtracts the Met residue (131.196 Da), and N-terminal acetylation adds $C_2H_2O$ (42.037 Da). The calculated mature masses are 28,103.3 Da for EGFP-Y40ONBN and 27,968.2 Da for EGFP-Y40N. ONBN differs from Asn by $C_7H_5NO_2$ (135.120 Da). In the supplied live-cell Prism dataset, the dominant −UV and +UV peaks occur at 28,102 and 27,974 Da, respectively, on a 2 Da mass grid. The estimated intact-mass tolerance was approximately ±5 Da for each spectrum. Using a conservative propagated bound of ±10 Da for the difference of two masses, the observed 128 Da decrease was consistent with the calculated 135.120 Da cage loss within this tolerance. Each plotted spectrum was normalized separately as $(I - I_{min})/(I_{max}$

− $I_{min}$), giving a maximum intensity of 1; normalized peak heights were not used to infer absolute protein amounts or decaging yield. For bacterial sfGFP, the supplied 247-residue coding sequence includes the C-terminal GSELHHHHH extension. After initiator-Met removal and chromophore maturation, the calculated average masses are 27,836.2 Da for D190ONBN and 27,701.1 Da for D190N; no bacterial N-terminal acetylation was assumed. The dominant unirradiated peak is 27,835 Da and the dominant 180 s peak is 27,703 Da. The expression construct was generated by replacing the Asp190 codon in the supplied backbone with TAG; the calculated sample masses therefore substitute ONBN or Asn at that position.

For peptide-level LC–MS/MS, purified bacterial sfGFP-D190ONBN expressed with ONBNRS_2 (Method 8) was digested with trypsin. Protein samples were denatured in 8 M urea and diluted with seven volumes of 50 mM Tris–HCl (pH 8.5) before addition of trypsin at an enzyme:protein ratio of 1:20 (w/w). Digestion proceeded at 37 °C for 16 h. Peptides were desalted using C18 tips (Thermo Fisher Scientific, 87784) and dried in a vacuum concentrator.

Peptides were separated on an Easy-nLC HPLC system coupled to an LTQ Orbitrap Elite mass spectrometer (Thermo Fisher Scientific). Solvent A was 0.1% formic acid in water, and solvent B was 0.1% formic acid in acetonitrile. The flow rate was 300 nL $min^{-1}$. The 70 min gradient used 3% B at 0 min, 10% at 10 min, 37% at 58 min, 100% at 59–64 min and 3% at 65–70 min. Full MS scans were collected in positive-ion mode over m/z 300–1,600 at a resolution of 120,000, with data-dependent MS/MS of the 15 most abundant precursor ions.

Raw spectra were processed using pParse 2.0 and searched with pFind 3.2.3. Searches used precursor and fragment-ion mass tolerances of 20 ppm, trypsin cleavage at the C terminus of Lys/Arg, and up to three missed cleavages. Carbamidomethylation of Cys and protein N-terminal acetylation were configured as fixed modifications; Met oxidation and ONBN were variable modifications. The peptide-spectrum-match false-discovery-rate threshold was 1%. The representative spectrum in Figure S8 has precursor charge 5+, m/z 924.85493 and a reported precursor mass error of +1.469 ppm; the original peak intensities and software ion annotations were retained.

##### 10. Mammalian cell culture and transfection

HEK293T cells were maintained as described in General Information and transfected at 70–90% confluency using Lipo8000 (Beyotime, C0533). Unless otherwise specified, transfections were performed in 12-well plates with 1–2 µg total DNA and 1.6 µL Lipo8000 per well. DNA and reagent were diluted separately in equal portions of a total of 50 µL Opti-MEM, combined and incubated for 10–15 min at room temperature before addition to cells. Medium was exchanged after 12–24 h, and expression proceeded for 48 h. The 10 cm dish preparation for cellular EGFP MS is specified separately in Method 14.

For amber suppression, the ONBNRS/tRNA and POI-TAG plasmids were cotransfected at a 1:1 DNA mass ratio unless stated otherwise, with or without 1 mM ONBN. Empty-vector controls used pCMV-Vector-A-4x7sk for most mammalian constructs and pCMV-Vector-B for FLuc and RLuc. ONBNRS_1 was used for RLuc, CAGE-prox$^{N}$/FLuc, glycosylation and cellular EGFP photodecaging MS; ONBNRS_2 was used for E2 experiments. Both variants were compared in the cellular EGFP fluorescence assay. Cell collection and immunoblot-lysate preparation followed Method 12. For luciferase activity measurements, parallel samples were lysed in the respective RLuc or FLuc kit buffer.

##### 11. UV irradiation and ONBN cellular compatibility

Photodecaging was performed with a 365 nm source. In the standard live-cell workflow, growth medium was replaced with PBS covering the cell monolayer, the plate was kept in an ice-water bath and the lid was removed during irradiation. A UVGO 365 nm curing lamp (80 W, AC 90–240 V) was positioned approximately 2 cm above the plate. Cells were irradiated for 2 min unless stated otherwise, corresponding to an incident dose of approximately 42 J $cm^{-2}$ at 350 mW $cm^{-2}$. PBS was removed after irradiation. Cells were either harvested immediately, as in the RLuc assay, or supplied with fresh growth medium for the designated postirradiation interval. Standard cell assays used 12-well plates; cellular EGFP MS used 10 cm dishes. Irradiation duration and the interval before harvest or readout are specified separately for each assay.

RLuc, E2 and glycosylation experiments used 2 min live-cell irradiation. RLuc cells were harvested immediately afterward. For live-cell EGFP MS, cells were irradiated for 2 min and harvested immediately, followed by Myc enrichment. The FLuc candidate screen irradiated lysates for 2 min. The detailed A348 activity series irradiated intact cells for 0, 10, 20, 30, 60, 90, 120 or 180 s, followed by immediate harvest and a lysate assay. A348 live-cell imaging and immunoblot series used the indicated 0, 30, 60, 90, 120 and 180 s exposures. Dark controls underwent the same handling without illumination.

HEK293T cells were seeded in 96-well plates at approximately 5,000 cells in 100 μL medium per well. ONBN was added at 0, 0.25, 0.5, 0.75, 1, 1.5, 2, 3 or 5 mM. No transfection or UV irradiation was performed. MTT measurements followed the Elabscience E-CK-A341 protocol: the 5× MTT reagent was diluted fivefold with the supplied diluent, 50 μL 1× MTT working solution was added per well, and cells were incubated for 4 h under culture conditions. Formazan Dissolution Buffer (100 μL) was added to each well and incubated for 1 h before endpoint absorbance was measured at 570 nm on the Synergy H1 reader. The plate protocol used a 37 °C temperature setpoint with preheating, followed by 30 s linear shaking at 360 cycles $min^{-1}$ (6 mm amplitude); the read used a 100 ms delay and eight measurements per data point. The dataset includes measurements at 0, 24, 48 and 72 h; Figure S10a presents the 48 h treatment-well $OD_{570}$ values, and Figure S10b presents the 0, 24, 48 and 72 h time courses. Control wells were measured separately; treatment-well absorbance is reported in raw OD570 units. Six treatment wells were measured per concentration. Exactly one highest and one lowest value were excluded from each concentration-by-time group, leaving four wells summarized as mean ± SD. The retained wells are within-assay replicates.

#### 12. General immunoblotting and source-image preparation

For routine immunoblotting, cells from each well of a 12-well plate were lysed on ice for 30 min in 40 μL Western and IP lysis buffer (Beyotime, P0013) containing 1× protease inhibitor cocktail (Beyotime; 100× stock diluted 1:100). Lysates were clarified at 12,000 × g for 10 min at 4 °C, and the supernatant was mixed with 10 μL 5× CWBIO denaturing reducing loading buffer (CW0027S) and heated at 95 °C for 10 min. Samples were prepared on a culture-well basis without BCA protein quantification. The PNGase F preparation used the lysis volume in Method 21. Proteins were separated on SurePAGE 4–20% gels (GenScript; M00655 or M00657 depending on well number), except that glycosylation reporters used 12% gels (GenScript, M00669), in 1× MOPS running buffer prepared from the GenScript 20× stock (M00680-500). The routine program was 90 V for 10 min followed by 160 V for 40 min. Vazyme or Thermo Fisher Scientific standards were run alongside samples. Immunoblot experiments were performed independently three times, and one representative set is shown for each experiment.

Proteins were transferred to methanol-activated 0.2 μm PVDF membranes using NCM Biotech rapid transfer buffer. The routine wet-transfer condition was 400 mA per transfer tank for 30 min at room temperature. Membranes were blocked with QuickBlock (Beyotime, P0252-500ml) for 10–15 min and incubated with CST monoclonal primary antibodies (1:1,000) overnight at 4 °C or for 1 h at room temperature. Antibodies and their catalog numbers are listed in the antibody table.

Membranes were washed three times with TBST for 5 min each after primary-antibody incubation, incubated with the appropriate CST anti-mouse or anti-rabbit secondary antibody (1:10,000) for 35–45 min at room temperature, and washed three times with TBST for 5 min each. Chemiluminescence was developed using equal volumes of the two Immobilon Western substrate components (Millipore, WBKLS0500) and recorded on the ChemiDoc system using Auto Optimal exposure unless otherwise specified. The same membrane was generally stripped with CWBIO stripping buffer and reprobed sequentially to detect the target proteins and loading controls. Any membrane cutting or nonadjacent-lane assembly is identified in the corresponding figure.

#### 13. EGFP fluorescence measurements and immunoblot validation of incorporation

HEK293T cells expressing EGFP-Y40TAG with ONBNRS_1 or ONBNRS_2 were cultured with or without ONBN. Fluorescence images were acquired using the Leica DMi8, an N PLAN 10×/0.25 dry objective, a GFP filter cube and a DFC420C camera controlled with LAS X 3.3.0.16799. Acquisition settings were held constant: exposure 423.525 ms, gain 1.8, gamma 0.9 and 3 × 3 binning, yielding 864 × 648-pixel images at 1.516 μm per pixel. The additional fields are presented in Figure S6a. Scale bars in Figure 1h correspond to 200 μm; the original scale rulers are retained in the supplementary fields.

Matched cell samples were harvested for immunoblotting. EGFP expression was detected using anti-Myc, and GAPDH was used as the loading control. Five numbered fields per group are displayed in Figure S6a. Images are presented as qualitative expression controls without image-based quantification. The displayed fields are technical subsamples from the same culture.

For cellular EGFP fluorescence measurements, HEK293T cells in 12-well plates were cotransfected with EGFP-Y40TAG and either ONBNRS_1/tRNA or ONBNRS_2/tRNA at a 1:1 DNA mass ratio under the conditions in Method 10. Cells were cultured for 48 h with or without 1 mM ONBN. Cells from each culture well were detached by trypsinization and washed once with PBS. Each sample, corresponding to one well of a 12-well plate, was transferred to one well of a conical-bottom 96-well plate and centrifuged at 500 × g for 15 min at room temperature. The supernatant was discarded, and pellet fluorescence was measured on the Synergy H1 reader at 488 nm excitation and 550 nm emission. Three culture wells were measured per condition; values are reported as mean ± SD (Figure S6b).

#### 14. Mammalian EGFP expression enrichment and live-cell photodecaging

Myc-tagged EGFP-Y40ONBN was expressed in HEK293T cells by cotransfecting pCMV-EGFP-Y40TAG with the ONBNRS_1/tRNA vector at a 1:1 DNA mass ratio using Lipo8000. Cells were cultured with 1 mM ONBN for 48 h. One 10 cm dish was used for each of the −UV and +UV conditions. The 10 cm dish transfection used 15 μg total DNA and 24 μL Lipo8000 in 10 mL medium, with 750 μL Opti-MEM used to prepare the mixture. Intact cells in the +UV dish were irradiated at 365 nm for 2 min and harvested immediately; the matched −UV dish underwent the same handling without irradiation.

Each dish was lysed in 1 mL Western and IP lysis buffer (Beyotime) containing 1× protease inhibitor cocktail freshly diluted 1:100 from the 100× stock. Lysates were clarified at 4,000 × g for 15 min at 4 °C. Each supernatant was incubated with 20 μL Myc-tag magnetic bead slurry (Beyotime) overnight at 4 °C. Beads were prewashed twice with TBST and washed three times with cold TBST after enrichment. Bound EGFP was eluted into 50 μL Myc peptide solution (150 μg $mL^{-1}$, Beyotime) for 2 h at 4 °C. Beads were magnetically separated and the peptide eluates were submitted directly for intact-protein UPLC–MS without ultrafiltration, concentration or solvent exchange.

#### 15. Optical activation and luminescence measurement of *Renilla* luciferase

HEK293T cells in 12-well plates were transfected with WT RLuc, RLuc-N53A or RLuc-N53TAG reporter plasmids at a WT:N53A:N53TAG DNA mass ratio of 1:2:5 across the three expression conditions. This ratio was chosen to reduce the large difference in expression between WT RLuc and the TAG construct before lysate adjustment. RLuc-N53TAG was coexpressed with ONBNRS_1/tRNA; cells were cultured with 1 mM ONBN for 48 h. Transfection used the standard Lipo8000/Opti-MEM procedure in Method 10. Mock-transfected cells served as the reference control; pCMV-Vector-B was the corresponding empty vector.

Intact cells were treated with or without 2 min UV irradiation and harvested immediately. For activity measurements, cells from each well of a 12-well plate were lysed in 100 μL reporter-gene lysis buffer supplied in the *Renilla* Luciferase Reporter Gene Assay Kit (Beyotime, RG016). Immunoblotting was used to select lysate-input proportions that gave approximately matched RLuc abundance for WT, N53A and N53ONBN in both UV conditions. WT and N53A lysates were diluted 20-fold and 40-fold, respectively, in the RLuc kit lysis buffer; N53ONBN and Mock lysates were used undiluted (WT:N53A:N53ONBN dilution factors, 20:40:1).

The same adjusted inputs were used for Figure 2f,g. Each assay contained 10 µL prepared lysate and 50 µL coelenterazine substrate working solution in an opaque 96-well plate. The supplied 100× coelenterazine stock was diluted 1:100 in assay buffer immediately before use.

The expression-adjusted RLuc assay was read on a Synergy H1 reader using Gen5 3.11.19, top optics, unfiltered emission, gain 200, a 1 s integration time, a 100 ms delay, extended dynamic range and a 4.5 mm read height. Three replicate measurements were used per UV condition. Values in Figure 2f are measured luminescence from the expression-adjusted lysates, summarized as mean ± SD. WT and N53A asterisks denote lysate-input adjustment. The separate unadjusted experiment in Figure S11 used gain 180 and the same integration time, optics, delay and read height; its first plate read was retained. Both datasets and their summary statistics are provided in Table S3F.

#### 16. Sequence alignment and structural analysis of E2 enzymes

Full-length human E2 amino-acid sequences were obtained from UniProt: UBE2A (P49459), UBE2C (O00762), UBE2D1 (P51668), UBE2D2 (P62837), UBE2D3 (P61077), UBE2E1 (P51965), UBE2N (P61088) and UBE2S (Q16763). All eight input sequences were verified against the corresponding human canonical entries. Multiple sequence alignment was performed in SnapGene using Clustal Omega. The saved alignment records specify FASTA input, CLUSTAL output and a line width of 60 residues. The conserved asparagine-containing active-site region was displayed while retaining full-length residue numbering. Human cofactor names follow UBE2V1 (Q13404) and UBE2V2 (Q15819).

Structural views were generated in PyMOL 3.1.6.1 from PDB 2GMI (chains A, B and C) and PDB 1J7D (chain B). The yeast Ubc13 chain in 2GMI carries C87S and is ester-linked to donor Ub. Figure 3c shows this complex with Ubc13, Mms2 and Ub in blue, purple and red, respectively, and N79 highlighted by a cyan semitransparent molecular surface. For Figure 3d, human WT UBE2N from 1J7D was rigidly superposed on 2GMI chain A using PyMOL align with Cα atoms and default outlier rejection (130 retained Cα pairs; RMSD 0.514 Å). The deposited Ub coordinates were retained. Human N79 and C87 and donor Ub-G76 were displayed as heavy-atom sticks, with their corresponding cartoon segments hidden. Ub-G76 is residue 576 in the deposited 2GMI coordinates. Green and blue dashed guides mark the N79 ND2–Ub-G76 O and C87 Sγ–Ub-G76 C separations in this rigid superposition, 5.007 and 1.021 Å, respectively. Rendering retained the deposited coordinates and the rigid alignment.

#### 17. Optical activation of UBE2N and analysis of TRAF6 ubiquitination

HEK293T cells in 12-well plates were cotransfected with ONBNRS_2/tRNA and a P2A-linked construct encoding Flag-tagged UBE2N-N79TAG, UBE2V1 (Uev1a) and HA-tagged TRAF6. The synthetase and E2-cassette plasmids were used at a 1:1 DNA mass ratio, with 0.5 µg of each plasmid per well. Expression proceeded for 48 h with 1 mM ONBN, using the standard 1.6 µL Lipo8000 and 50 µL Opti-MEM mixture. Samples without ONBN assessed amber-suppression-dependent UBE2N expression.

For the main time course in Figure 3g, cells expressing UBE2N-N79ONBN and UBE2V1 were irradiated for 2 min and harvested at 0.5, 1, 1.5 and 2 h after irradiation. A nonirradiated control was analyzed alongside the time course. WT UBE2N and UBE2N-N79A were tested with UBE2V1 and TRAF6 using the same collection series (Figure S12c). ONBN-containing UBE2N cofactor comparisons using UBE2V1 (Uev1a) or UBE2V2 (Mms2) are shown in Figure S13.

TRAF6 ubiquitination-associated bands were examined directly in cell lysates prepared with Western and IP lysis buffer and 1× protease inhibitor cocktail (Beyotime), without affinity enrichment. HA-tagged TRAF6 and its high-molecular-weight species were detected with anti-HA. Flag-tagged UBE2N and GAPDH were detected in the corresponding samples. The increase in high-molecular-weight TRAF6-reactive signal was assessed qualitatively; these blots were not subjected to densitometric quantification or ubiquitin-chain linkage assignment.

For the nonreducing UBE2N–Ub assay (Figure S12b), cells expressing Flag-tagged UBE2N-N79ONBN with Ub were lysed in Western and IP lysis buffer containing 1× protease inhibitor cocktail. Lysates were mixed with 4× NuPAGE LDS Sample Buffer (Thermo Fisher Scientific, NP0007) to 1× without added reducing reagent and heated at 70 °C for 20 min. The +UV sample was harvested 2 h after 2 min irradiation. Samples were resolved on 4–20% SurePAGE gels in 1× MOPS running buffer using 90 V for 10 min followed by 160 V for 40 min; transfer and anti-Flag/GAPDH detection followed Method 12. The upper UBE2N-associated Ub band was evaluated qualitatively.

#### 18. FLuc complex modeling and CAGE-prox$^{N}$ site selection

##### 18.1 Modeling and relaxation of the FLuc–luciferyl-AMP complex

The FLuc–luciferyl-AMP model (Figure S14) was prepared for Rosetta by adding hydrogens and parameterizing the ligand (adenylated D-luciferin). Following the CAGE-prox workflow[7], the complex underwent restrained torsional relaxation (40 models), followed by Cartesian relaxation with ramped restraints (200 models) in Rosetta[8]. The lowest-scoring model from each stage was retained. The final model (Figure 4b) contained 550 FLuc residues in chain B and one luciferyl-AMP molecule in chain X. Structures were visualized in PyMOL.

##### 18.2 Geometric and energetic evaluation of FLuc sites

Geometric screening used the final relaxed FLuc–luciferyl-AMP model shown in Figure 4b, with all 550 FLuc residues in chain B and ligand atoms in chain X. The distance is the shortest Euclidean atom-center distance between any existing atom of a residue, including backbone, side-chain and hydrogen atoms, and any existing ligand atom, including hydrogen atoms. For a non-Gly residue, the angle is the smallest angle between the vector from Cα to Cβ and any vector from Cα to a ligand heavy atom. Angles are reported in degrees. Gly is assigned an angle of 0° by convention because it lacks Cβ. The distance and angle minima may involve different ligand atoms. The modified CAGE-prox criteria[7], a distance below 6.5 Å and an angle below 120°, retained 59 of the 550 residue positions. All 59 had Rosetta energy records.

Each geometrically selected position was evaluated for substitution with Asn using the Rosetta ref2015_cart score function[8]. Three rounds of optimization were performed for each wild-type and mutant structure. The original energy scores are provided in Table S2A.

The folding-stability score change, $\Delta G_f$, was calculated as the minimum mutant score minus the minimum wild-type score. The folding cutoff was $\Delta G_f < 0.2$ Rosetta energy units (REU), retaining ten positions: N197, S199, G200, N229, G246, T251, G339, A348, D422 and K529. Figure 4d plots these scores against FLuc residue number on linear axes; the dotted line marks the 0.2 REU cutoff. Energy scores are provided in Table S2A. Complete geometric values are in the CAGE-prox geometry worksheet of ONBN_Source_Data.xlsx.

The computational selection retained 59 of 550 residues by geometry and ten positions by the folding criterion. The final experimental screen includes these ten positions, N197, S199, G200, N229, G246, T251, G339, A348, D422 and K529, together with WT FLuc. Figure 4e therefore contains ten ONBN-incorporation variants and WT.

#### 19. Experimental evaluation of proximal FLuc sites and UV dose response

The ten FLuc positions listed in Method 18 were individually replaced with TAG for ONBN incorporation and compared with WT FLuc (Figure S15; Table S3C). HEK293T cells in 12-well plates were cotransfected with ONBNRS_1/tRNA and FLuc plasmids at a 1:1 DNA mass ratio using the standard conditions in Method 10. Expression proceeded for 48 h with 1 mM ONBN; pCMV-Vector-B was the empty-vector control. Following cell harvest, samples were lysed using the buffer supplied in the Firefly Luciferase Reporter Gene Assay Kit (Yeasen, 11401), with 70 µL lysate for each −UV or +UV sample. Lysates were irradiated for 2 min or kept unirradiated. Each sample was divided into three 10 µL aliquots, and each aliquot was mixed with 100 µL luciferin substrate working solution in an opaque 96-well plate. Luminescence was measured on the Synergy H1 reader in Gen5 3.11.19 using top optics, unfiltered emission, gain 200, a 1 s integration time, a 100 ms delay, extended dynamic range and a 4.5 mm read height. The read region was A1–F12. Results are shown as mean ± SEM of three measurements. Expression was assessed by anti-His immunoblotting, with GAPDH as the loading control (Figure S15c).

For each candidate and the WT reference, the activation ratio was mean(+UV luminescence)/mean(−UV luminescence), with three replicates per UV condition. Figure 4e uses mean(+UV luminescence)/10 on the horizontal axis and this untransformed activation ratio on the vertical axis; both axes are logarithmic. The horizontal axis uses a common 1/10 scaling of the mean +UV signal. Ratios were calculated from the original measurements, without blank subtraction, before applying logarithmic axis scales. In the final screen, A348 has $x$ = 327.2333 and $y$ = 29.7485; WT has $x$ = 34305.1 and $y$ = 1.07345. Figure S15 presents the original luminescence values as mean ± SEM ($n$ = 3 per condition) and the activation ratio with SEM propagated from the two group means. Numerical values are provided in Table S3C.

For the detailed FLuc-A348ONBN activity series, intact cells were irradiated for 0, 10, 20, 30, 60, 90, 120 or 180 s and harvested immediately. Cells were lysed in the FLuc kit buffer, and activity was measured using 10 µL lysate and 50 µL assay reagent. The detailed A348 assay used gain 200 with the same 1 s integration time, top optics, unfiltered emission, 100 ms delay, extended dynamic range and 4.5 mm read height. Each time point contained four measurements; results are shown as mean ± SEM. Figure 4f retains 0, 30, 60, 90, 120 and 180 s; Figure S16a includes all eight times and individual values, with source measurements in Table S3E. The site screen irradiated lysates for 2 min; the detailed A348 experiment irradiated intact cells before lysis. Separate cells were used for imaging in Figure 4g. For Figure 4h, parallel cells were irradiated for the indicated durations and harvested immediately at each time point for routine anti-His and GAPDH immunoblotting (Method 12). Photodecaging at A348 produces A348N.

For live-cell imaging, HEK293T cells were prepared in 3.5 cm dishes using the same transfection conditions as a 6-well culture: 2.5 µg total DNA at a 1:1 synthetase:FLuc mass ratio, 4 µL Lipo8000 and 125 µL total Opti-MEM in 2 mL culture medium per dish. Cells were cultured for 48 h with 1 mM ONBN and irradiated for the indicated durations. After irradiation, medium was replaced with 1 mL of 1 mM D-luciferin sodium salt in PBS, diluted from a 100 mM stock. Six dishes were imaged together in a single ChemiDoc chemiluminescence acquisition using Auto Rapid exposure. The original grayscale full-field export and the corresponding pseudocolor rendering are shown in Figure S16c,d. Six equal-diameter circular regions (1,300 pixels) were cropped within the dish interiors to exclude the walls. A common piecewise-linear blue–cyan–yellow–red lookup table was applied to the original 8-bit grayscale values using a shared linear display window of 0–140. Values above 140 were assigned the highest color (red) for display; the original 8-bit intensities were unchanged. The same mapping was used for the full field and every crop, without per-dish intensity normalization, background subtraction, spatial filtering or geometric stretching. Crop coordinates and the lookup table were retained with the source images. This imaging workflow followed the live-cell luminescence approach in the PABD study[6], with ONBN photodecaging replacing chemical activation. The relative-intensity color scale is shown alongside the full-field pseudocolor image. Expression controls were prepared by routine immunoblotting (Method 12).

#### 20. ER reporter design and time-resolved glycosylation

An ER-targeted EGFP reporter was engineered with an N-terminal signal peptide and a C-terminal glycosylation sequon, Flag tag, and KDEL sequence. Full reference coding sequences, including linkers, are provided in the Protein sequences section of Supplementary Data. The FANGT baseline and XCNGT series differ at the local sequon as listed in Table S1C. FANGT and WCNGT were selected from previously optimized enhanced aromatic sequons[9,10]. The XCNGT series varies the −2 residue around the WCNGT context to measure its effects on the glycosylation time course.

For the baseline reporter, the glycan-acceptor Asn in FANGT was replaced with TAG and expressed as FA-ONBN-GT. HEK293T cells in 12-well plates were cotransfected with the reporter and ONBNRS_1/tRNA plasmids at a 1:1 DNA mass ratio, using 1–2 µg total DNA, 1.6 µL Lipo8000 and 50 µL total Opti-MEM per well. Cells were cultured with 1 mM ONBN for 48 h. Medium was replaced with ONBN-free medium before 2 min live-cell UV irradiation. The displayed sampling series comprised 0, 0.5, 1, 1.5, 2 and 3 h, with 2 min UV irradiation where indicated. The XCNGT variants used the same transfection and collection procedure. The time courses are displayed from 0 to 3 h.

Cells were lysed in Western and IP lysis buffer (Beyotime) and prepared for reducing SDS–PAGE according to Method 12. Glycosylation reporters were resolved on 12% SurePAGE gels (GenScript) and detected by anti-Flag immunoblotting, with GAPDH as the loading control. The upper reporter band was assigned as glycosylated EGFP by its sensitivity to PNGase F treatment.

##### 21. PNGase F validation of N-linked glycosylation

Cells expressing the FANGT- or WCNGT-derived reporter were harvested 3 h after UV irradiation and lysed using 50 μL Western and IP lysis buffer per well of a 12-well plate, with 1× protease inhibitor cocktail. Clarified lysate (9 μL) was mixed with 1 μL 10× Denaturing Buffer from the PNGase F Deglycosylation Kit (Beyotime, P2318S), heated at 100 °C for 10 min, cooled on ice and briefly centrifuged. Reaction Buffer, Renaturing Buffer and PNGase F were then added according to the manufacturer's denaturing protocol. Enzyme digestion was performed overnight at 37 °C. Mock-digested aliquots underwent the same buffer and heat treatment without enzyme.

Samples were analyzed by anti-Flag immunoblotting to compare the PNGase F-sensitive upper reporter band with the lower band. GAPDH was detected in the corresponding samples.

##### 22. Sequon-dependent kinetics and immunoblot densitometry

The −2 position was varied within a common XCNGT scaffold, retaining Cys at −1 and NGT as the acceptor sequon after decaging. The six variants in Figure 5d are ACNGT, FCNGT, VCNGT, CCNGT, RCNGT and DCNGT, in that row order. Figure S19 includes all nine XCNGT variants, with X = A, F, V, C, R, D, G, P and W. ONBN initially occupies the acceptor Asn position. The FANGT baseline is analyzed separately because its −1 residue is Ala.

For each variant, the 0, 0.5, 1, 1.5, 2 and 3 h series were analyzed by immunoblotting. Images were processed using the Subtract Background command in Fiji (ImageJ), and glycosylated (G) and nonglycosylated (N) EGFP bands were measured as integrated density (IntDen) within consistently defined regions. G/N was calculated by dividing the background-subtracted integrated density of G by that of N. Values are summarized as mean ± SD of three measurements. Only unsaturated images were used.

Figure 5d displays the mean G/N values for the six main-series variants on a common heatmap scale of 0–0.3. The displayed columns are 0, 0.5, 1, 1.5, 2 and 3 h. G/N is the glycosylated-band intensity divided by the nonglycosylated-band intensity. The heatmap uses the original G/N values on one common scale. Figure S19 provides immunoblots and the complete G/N time courses for all nine XCNGT variants. Integrated band intensities and G/N measurements are in the XCNGT worksheet of ONBN_Source_Data.xlsx.

#### Supplementary Figures

##### Figure S1

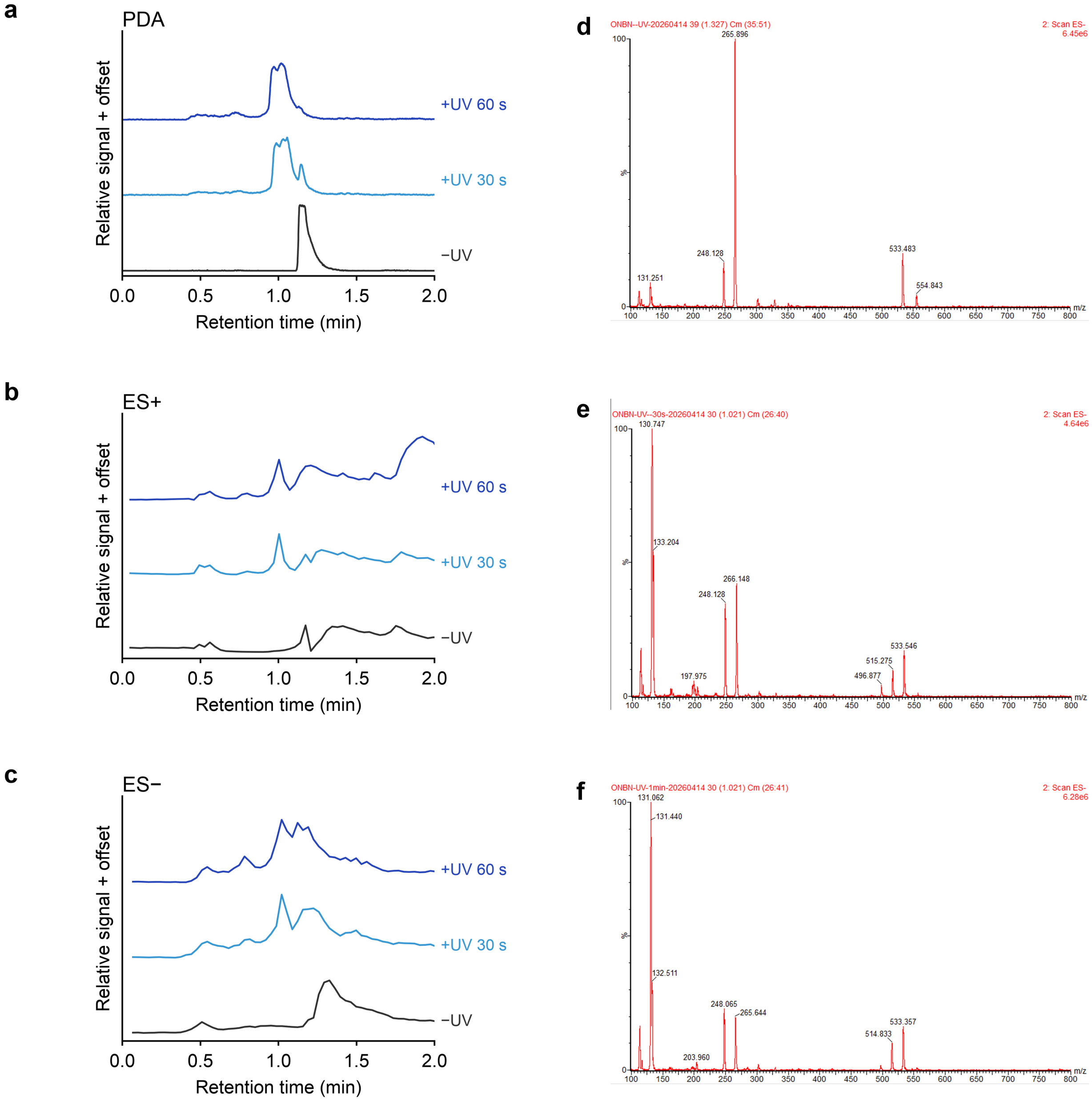


**Figure S1.** Small-molecule photodecaging of ONBN. (a–c) PDA, positive-ion and negative-ion chromatograms, respectively, before irradiation and after 30 or 60 s irradiation at 365 nm. The 0–2 min retention-time window is shown. Within each channel, all three traces retain one common normalization and vertical offsets. (d–f) Original negative-ion mass spectra extracted around the corresponding chromatographic peaks for the dark, 30 s and 60 s samples, respectively. Native peak shapes, intensities and m/z annotations are retained. The stronger signal near nominal m/z 131 after irradiation is consistent with [Asn−H]−; the ONBN-associated ion is near m/z 266.

#### Figure S2

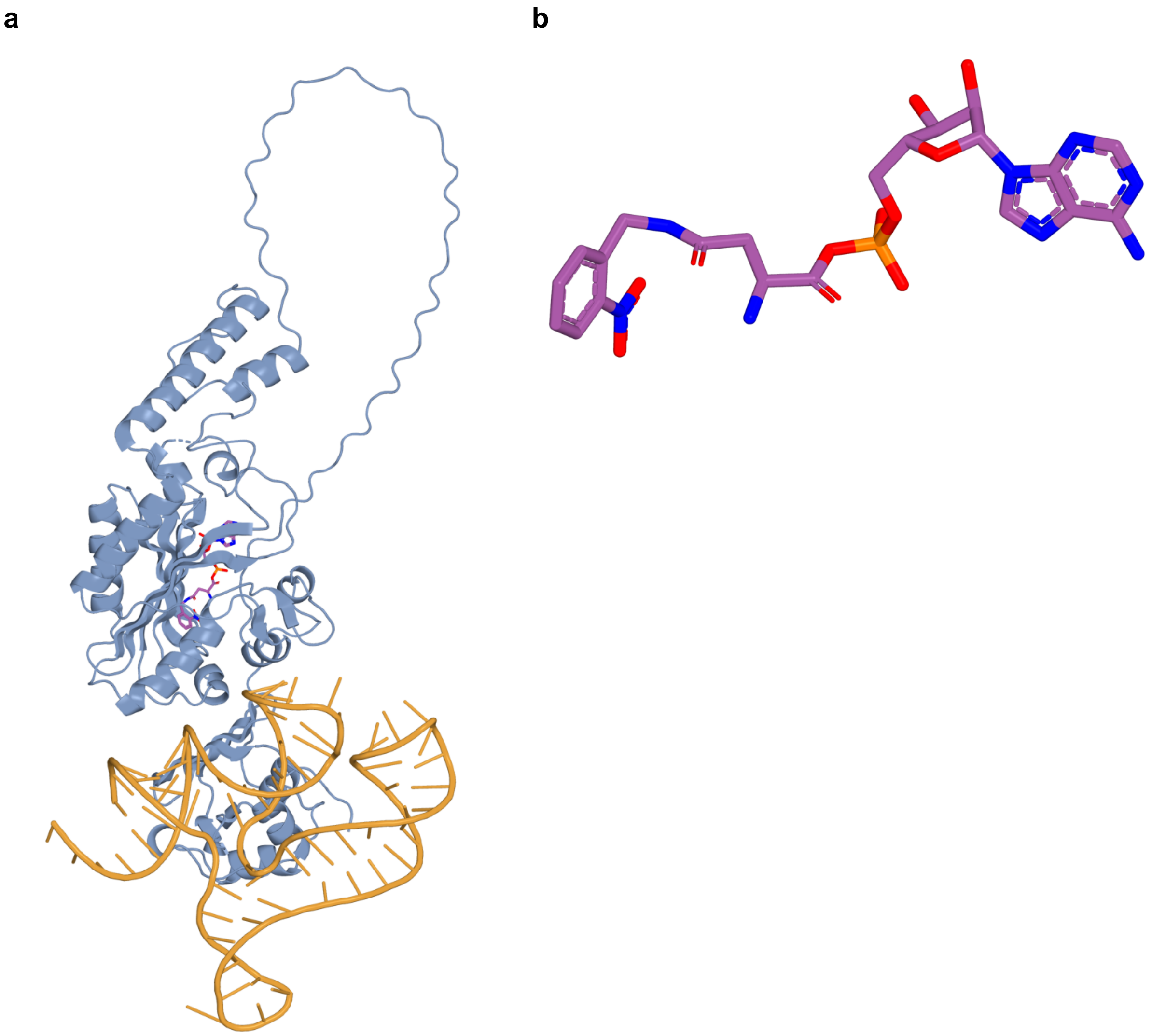


**Figure S2.** Predicted BnDRS–PylT–ONBN–AMP complex. (a) Overall view of the saved model, with BnDRS in slate blue, the suppressor tRNA in gold and ONBN–AMP shown as sticks. (b) ONBN–AMP conformation in the same model. Ligand carbon atoms are purple; nitrogen, oxygen, sulfur and phosphorus follow blue, red, yellow and orange coloring. All views were generated from the supplied PyMOL session without coordinate optimization or addition of a comparison structure.

#### Figure S3

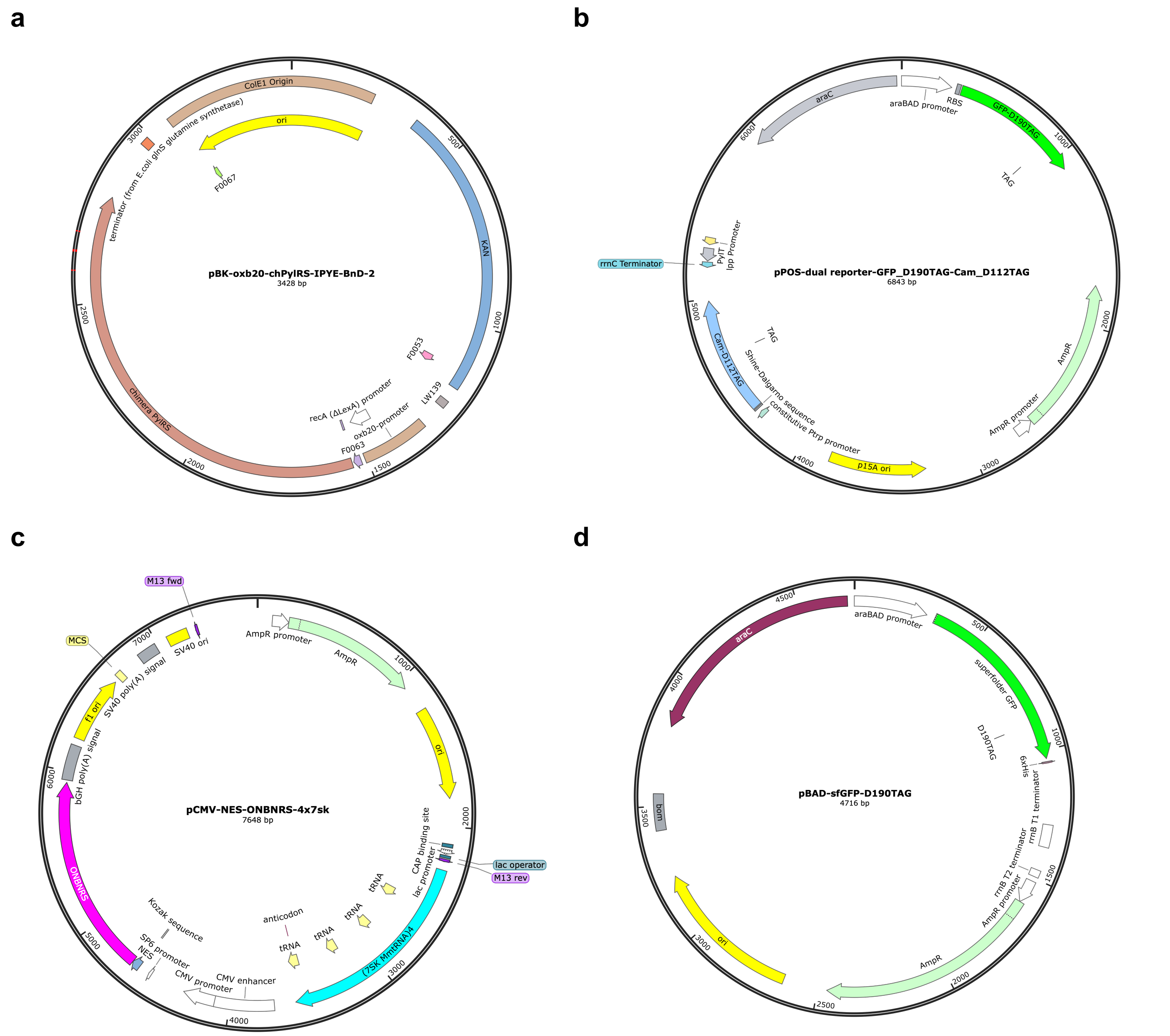


**Figure S3.** Plasmid maps for ONBNRS library construction, selection and expression assays. (a) Organization of the kanamycin-resistant pBK synthetase-library vector with an oxb20 promoter. The map was exported from pBK-oxb20-chPylRS-IPYE-BnD-2 as a backbone-layout reference; the library scaffold was BnDRS. (b) The ampicillin-resistant A1-pos-Cam-D112TAG-GFPuv-D190TAG-PylT reporter, carrying CAT-D112TAG, GFPuv-D190TAG with a C-terminal $His_6$ tag, and PylT. CAT expression is driven by a constitutive Ptrp promoter; GFPuv is induced from araBAD. The reporter map was exported directly from its SnapGene file. (c) Native SnapGene map of pRS (pCMV-NES-ONBNRS-4x7sk), showing the CMV-driven NES–ONBNRS_1 cassette, four 7SK–tRNA cassettes, ampicillin resistance and replication origin. The NES is fused to the synthetase N terminus. **(d)** The pBAD-sfGFP-D190TAG reporter used for bacterial incorporation assays.

**Figure S5**

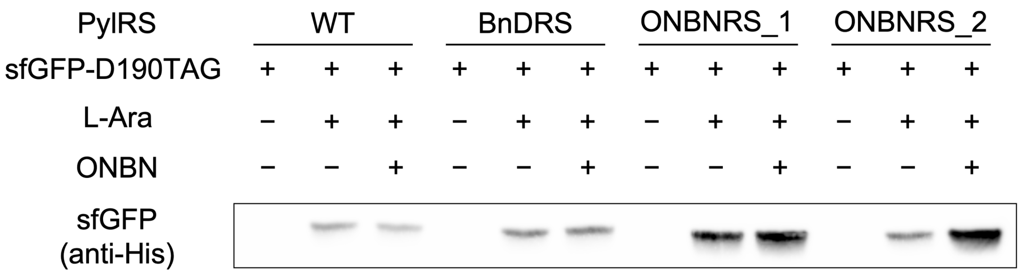


**Figure S5.** Immunoblot validation of bacterial ONBN incorporation. Anti-His immunoblot of sfGFP-D190TAG expressed in *E. coli* with WT chPylRS, BnDRS, ONBNRS_1 or ONBNRS_2 under the indicated L-arabinose and ONBN conditions. Corresponding fluorescence measurements are shown in Figure 1g. Immunoblots are representative of three independent experiments.

Figure S6

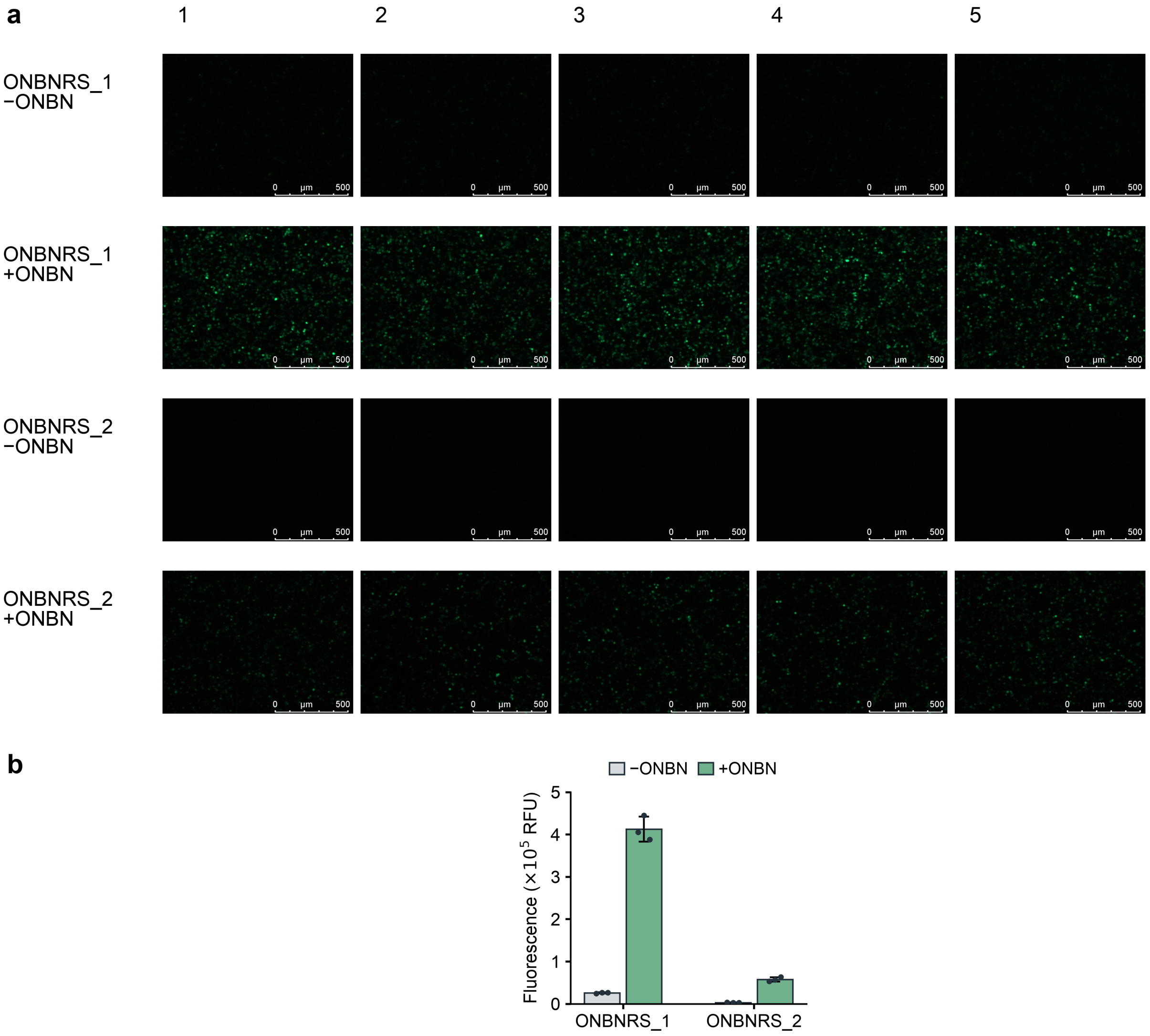


**Figure S6.** Cellular EGFP expression with ONBNRS variants. (a) HEK293T cells expressing EGFP-Y40TAG with ONBNRS_1 or ONBNRS_2, with or without ONBN. Five fields are shown per condition under identical Leica acquisition settings; original calibrated scale rulers are retained. (b) Plate-reader fluorescence for the four conditions shown in (a). Bars show mean ± SD; dots indicate individual culture wells ($n$ = 3 per condition). Images are qualitative expression controls; no image-based quantification was performed.

#### Figure S7

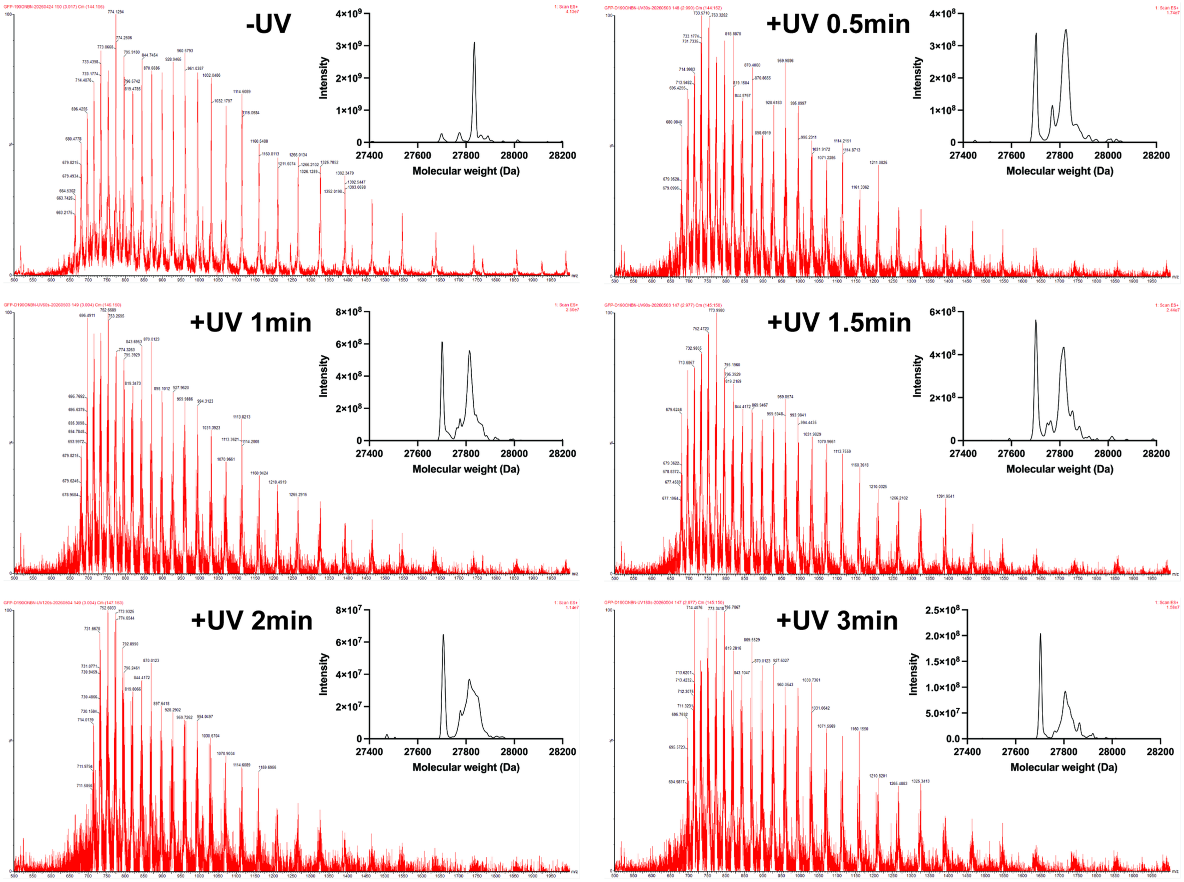


**Figure S7.** In vitro photodecaging of bacterial sfGFP-D190ONBN expressed with ONBNRS_2. The mass-spectrometry figure shows charge-state envelopes and the corresponding deconvoluted intact-protein spectra before irradiation and after 0.5, 1, 1.5, 2 and 3 min UV irradiation. Samples contained 50 µL purified protein at 0.5 mg $mL^{-1}$ in a 96-well plate kept in an ice-water bath and were analyzed directly by UPLC–MS. Calculated average masses after initiator-Met removal and chromophore maturation are 27,836.2 Da for D190ONBN and 27,701.1 Da for D190N. Dominant observed peaks are 27,835 Da before irradiation and 27,703 Da at 3 min.

#### Figure S8

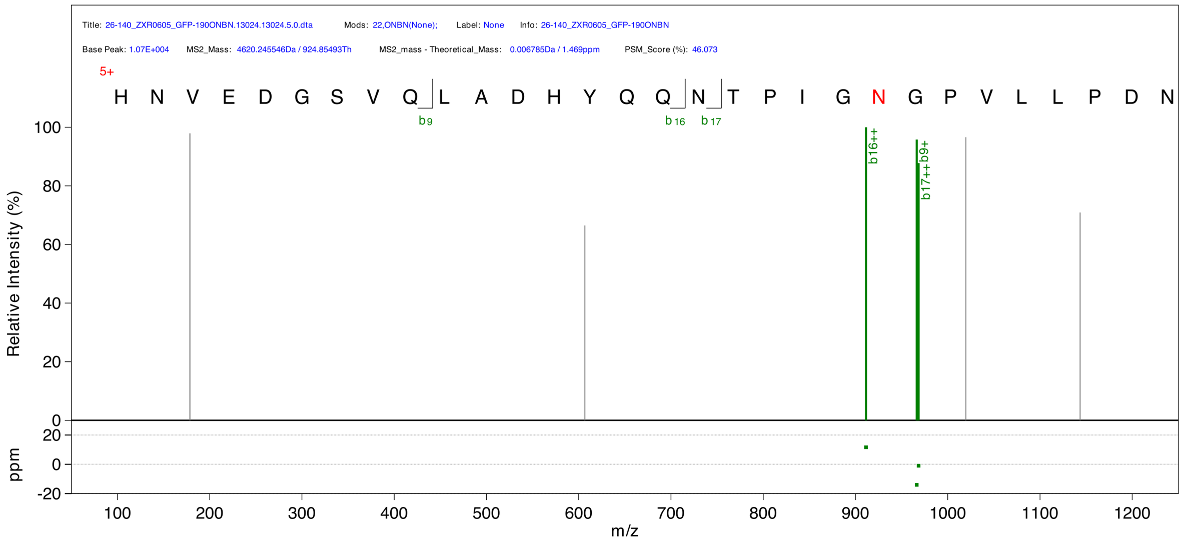


**Figure S8.** LC–MS/MS characterization of bacterial sfGFP-D190ONBN. Purified sfGFP-D190ONBN expressed with ONBNRS_2 was digested with trypsin. The displayed spectrum corresponds to the peptide HNVEDGSVQLADHYQQNTPIGXGPVLLPDNHYLSTQSVLSK, where X denotes ONBN at sfGFP residue 190 (peptide position 22). The software export marks this position with a red N and lists ONBN in the spectrum header. The precursor ion has charge 5+ and m/z 924.85493, with a reported mass error of +1.469 ppm. Green peaks and labels indicate software-assigned b ions; the lower panel shows fragment-ion mass errors in ppm. Peak positions, intensities and ion annotations were retained from the native export.

#### Figure S9

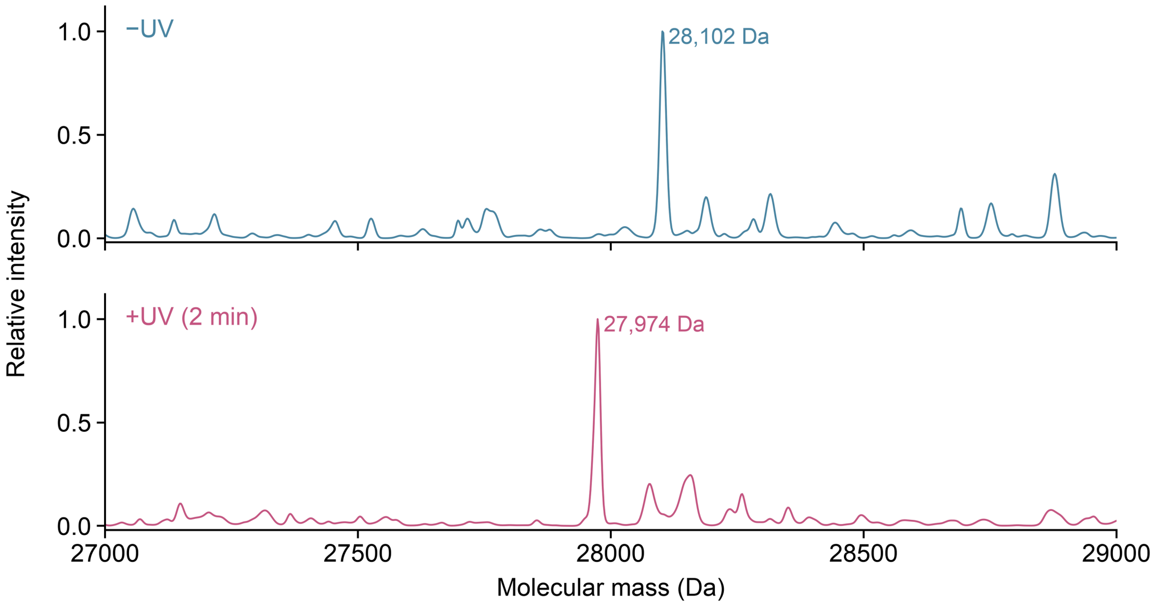


**Figure S9.** Photodecaging of EGFP-Y40ONBN in living cells. Myc-tagged EGFP-Y40ONBN was expressed with ONBNRS_1, irradiated as intact cells, harvested immediately and enriched using Myc beads. Theoretical average masses for the mature, initiator-Met-cleaved and N-terminally acetylated proteoforms are 28,103.3 Da (caged) and 27,968.2 Da (decaged); dominant observed peaks are 28,102 and 27,974 Da. The two spectra are shown separately on matched axes, each independently normalized to a maximum of 1 after minimum-intensity subtraction. The 128 Da observed decrease is consistent with the calculated 135.120 Da cage loss within a conservative ±10 Da difference tolerance derived from approximately ±5 Da per intact mass (Method 9).

#### Figure S10

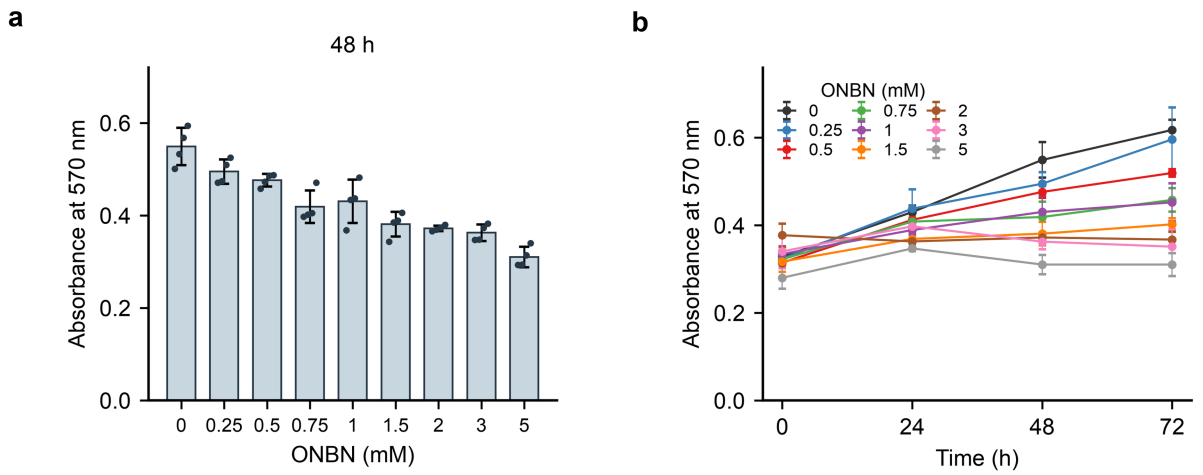


**Figure S10.** ONBN concentration and exposure-time dependence of the MTT readout. (a) Raw treatment-well absorbance at 570 nm after 48 h at the indicated ONBN concentrations. (b) Raw absorbance time courses at 0, 24, 48 and 72 h for the same nine concentrations. Cells received no transfection or UV irradiation. Control wells were measured, but values were not control-subtracted or normalized. At each concentration and time point, exactly one highest and one lowest value among six wells were excluded; bars or points show the mean and error bars show SD of the four retained values ($n$ = 4). Dots in a show retained values. Lines in b connect measured time points. Source readings are in the MTT worksheet of ONBN_Source_Data.xlsx.

**Figure S11**

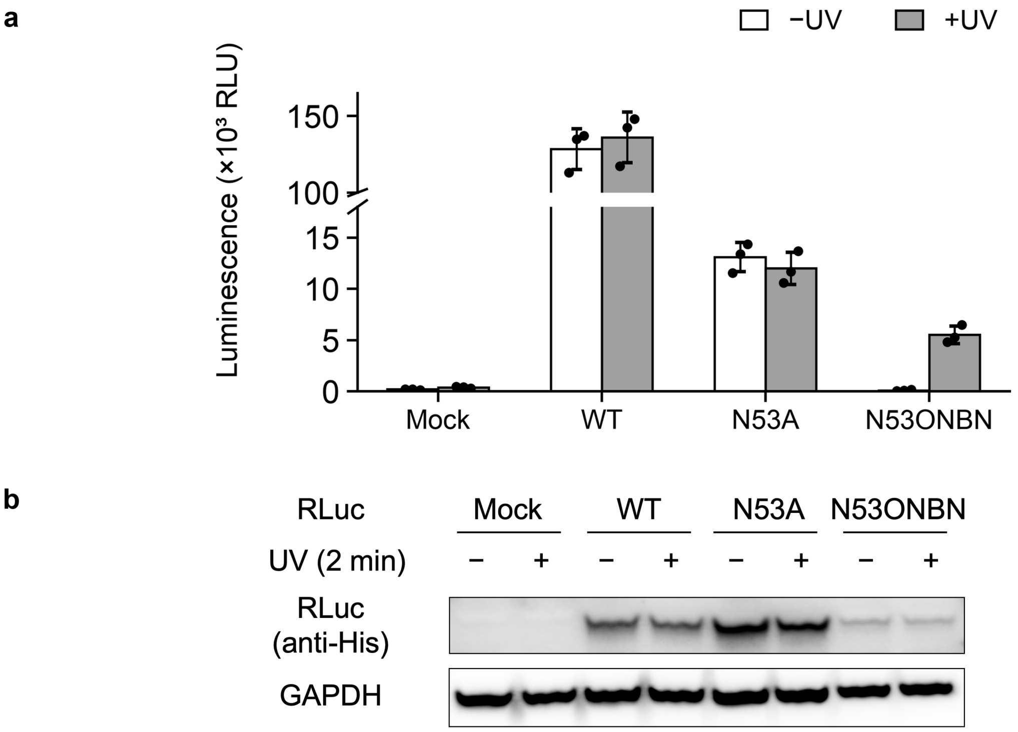


**Figure S11.** RLuc activity without protein-abundance adjustment and corresponding expression controls. (a) Raw luminescence for Mock, WT, N53A and N53ONBN samples with or without 2 min UV irradiation. RLuc-N53ONBN was expressed with ONBNRS_1; intact cells were irradiated and harvested immediately. Bars and error bars show mean ± SD ($n = 3$); dots show individual measurements from the first plate read. Values were not mock-subtracted or adjusted for protein abundance. The broken y axis omits 18,000–100,000 RLU (axis displayed in units of $10^3$ RLU). Numerical source data are provided in Table S3F. (b) Corresponding anti-His immunoblot of RLuc with GAPDH as the loading control. Lanes are Mock, WT, N53A and N53ONBN, each in −UV/+UV order. Immunoblots are representative of three independent experiments.

**Figure S12**

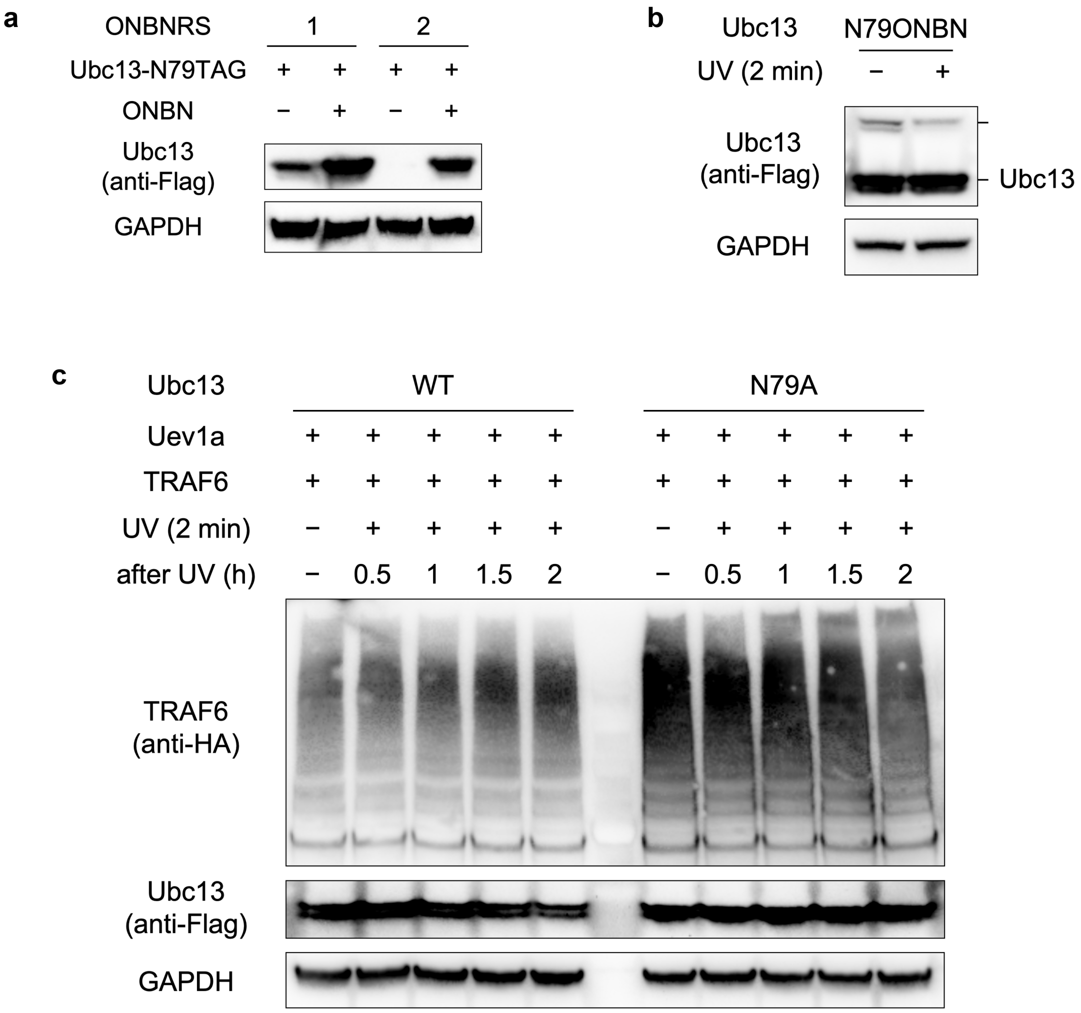


**Figure S12.** UBE2N expression and control assays. (a) ONBN-dependent expression of Flag-tagged UBE2N-N79TAG with ONBNRS_1 or ONBNRS_2. (b) Nonreducing anti-Flag immunoblot of UBE2N-N79ONBN coexpressed with Ub, with or without 2 min live-cell UV irradiation. The +UV sample was harvested 2 h after irradiation. (c) TRAF6 time courses with WT UBE2N or UBE2N-N79A coexpressed with UBE2V1 (Uev1a) and TRAF6. Lanes within each group are −UV followed by 0.5, 1, 1.5 and 2 h after 2 min irradiation. Anti-HA detects TRAF6, anti-Flag detects UBE2N and GAPDH is the loading control. Nonreducing preparation and electrophoresis conditions are given in Method 17. Immunoblots are representative of three independent experiments.

**Figure S13**

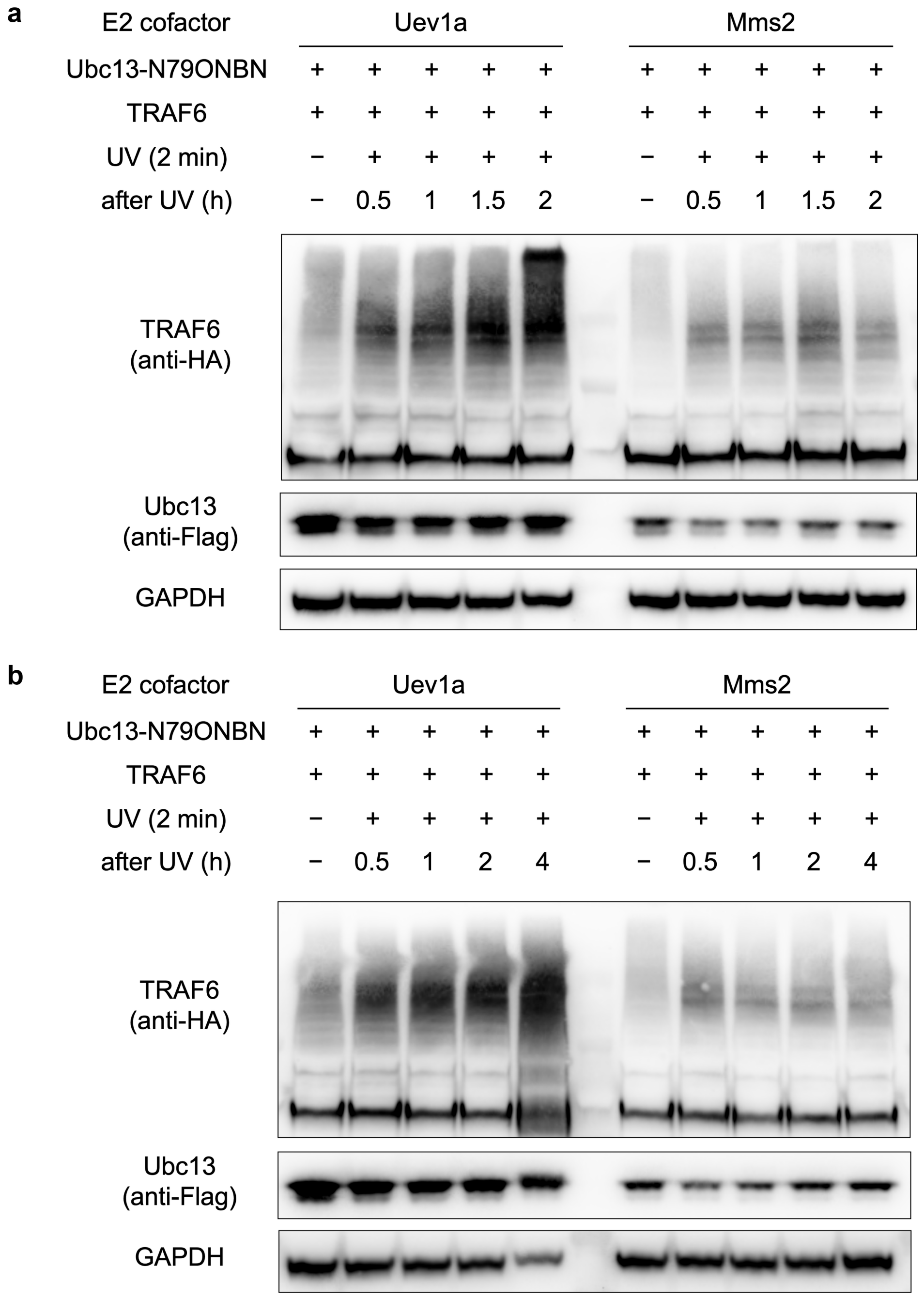


**Figure S13.** UBE2N cofactor comparisons. (a) TRAF6-reactive bands after ONBN-containing UBE2N was coexpressed with UBE2V1 (Uev1a) or UBE2V2 (Mms2), with collection at 0.5, 1, 1.5 and 2 h after 2 min UV irradiation. (b) Extended series collected at 0.5, 1, 2 and 4 h. Anti-HA detects TRAF6, anti-Flag detects UBE2N and GAPDH serves as the loading control. The high-molecular-weight bands are presented qualitatively. Immunoblots are representative of three independent experiments.

**Figure S14**

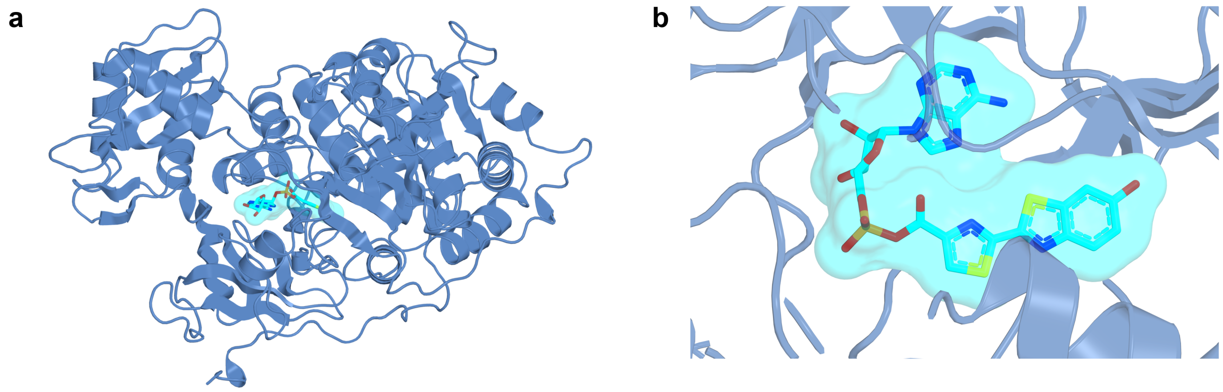


**Figure S14.** Predicted FLuc–luciferyl-AMP complex before relaxation. (a) Overall model. (b) Close-up of the ligand pocket in the same model. FLuc is shown as a blue cartoon; luciferyl-AMP is shown as sticks with a cyan semitransparent molecular surface. Nitrogen, oxygen, sulfur and phosphorus atoms are blue, red, yellow and orange, respectively. Views were generated from the pre-relaxation model without further coordinate optimization. The final relaxed model is used for the main-text structural view in Figure 4b (Method 18).

**Figure S15**

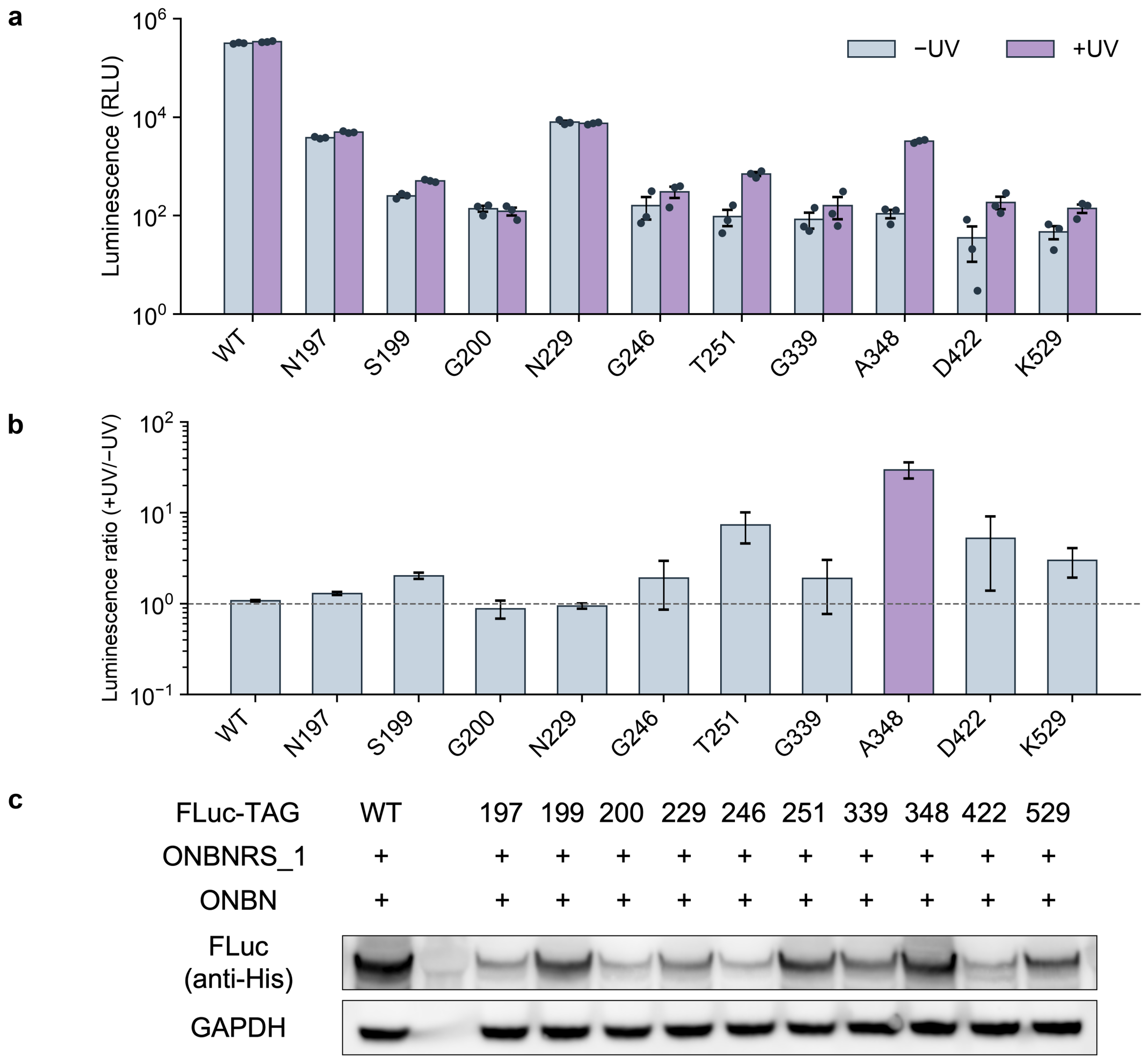


**Figure S15.** Activity screen of WT FLuc and 10 ONBN-incorporation variants. (a) Measured luminescence before and after 2 min irradiation of cell lysates, shown as individual values and mean ± SEM ($n = 3$). The ordinate is logarithmic. (b) +UV/−UV ratios calculated from group means, with propagated SEM. Source measurements and statistics are in Table S3C. (c) Anti-His immunoblots of WT FLuc and the ten indicated ONBN-incorporation variants, with GAPDH as the loading control.

#### Figure S16

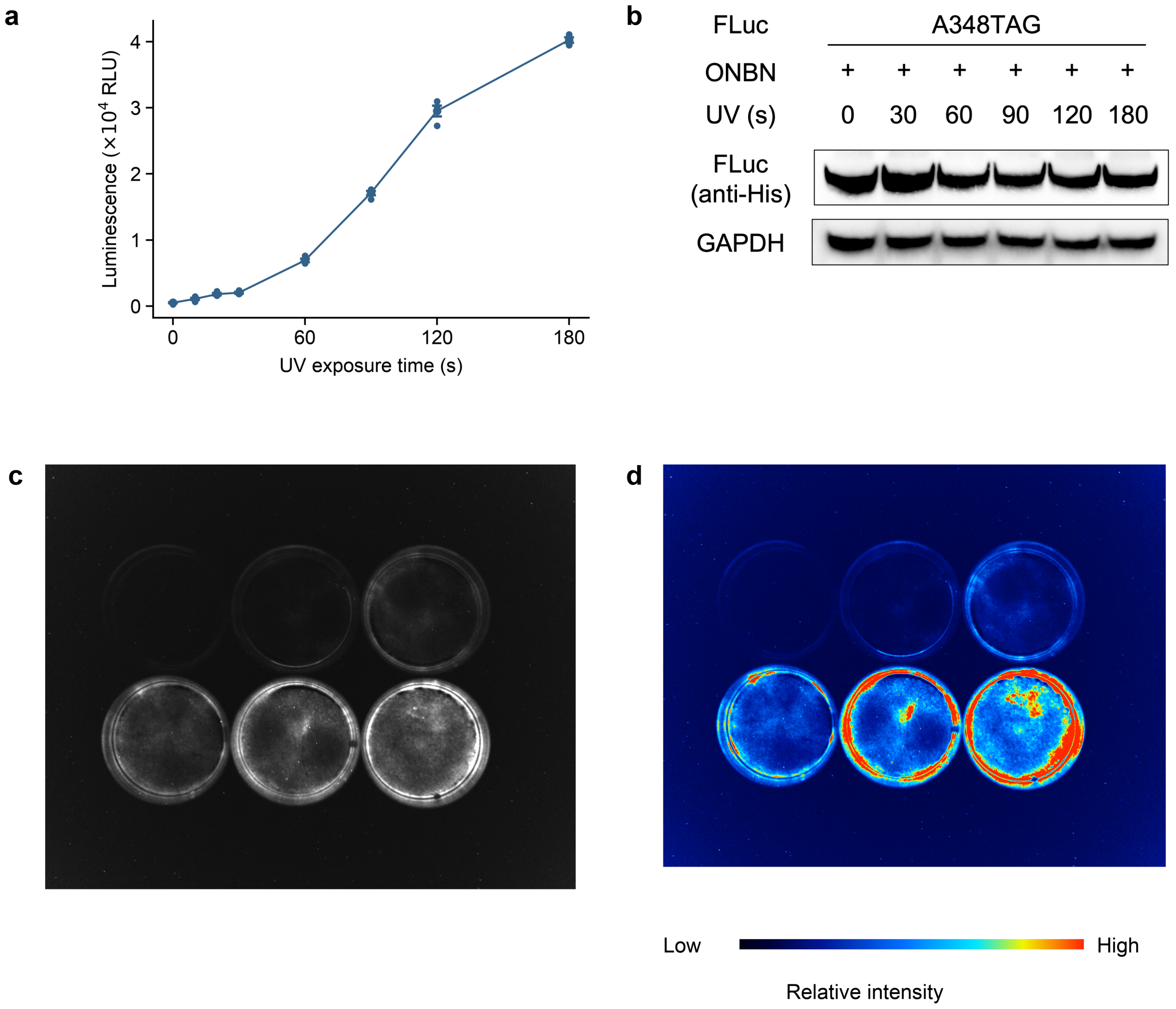


**Figure S16.** Detailed activation of FLuc-A348ONBN and full-field live-cell luminescence imaging. (a) Complete 0, 10, 20, 30, 60, 90, 120 and 180 s UV-exposure series. Intact cells were irradiated and harvested immediately for the lysate assay; individual background-subtracted measurements and mean ± SEM are shown ($n = 4$). Source values are in Table S3E. (b) Anti-His immunoblot after 0, 30, 60, 90, 120 and 180 s irradiation, with GAPDH as the loading control. (c) Original grayscale full-field acquisition of six dishes. (d) The same field displayed with the shared blue–cyan–yellow–red lookup table and 0–140 display window; values above 140 are red. The color scale indicates relative signal intensity. Upper row, 0, 30 and 60 s; lower row, 90, 120 and 180 s, from left to right. The main-text circular images originate from this field. No per-dish normalization or spatial filtering was applied. Immunoblots are representative of three independent experiments.

#### Figure S17

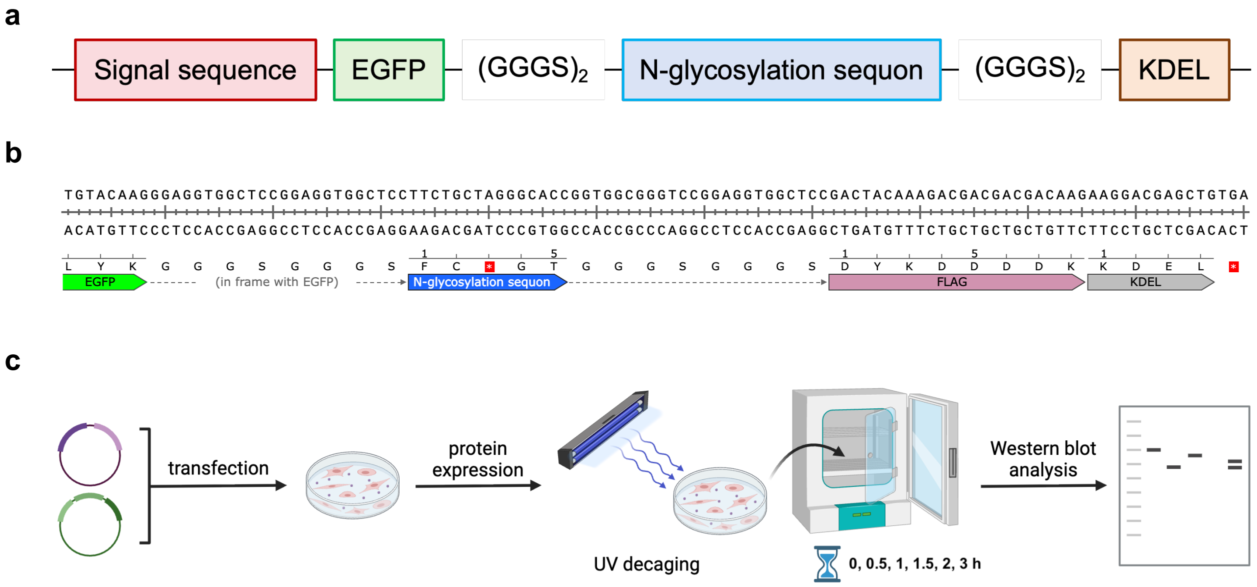


**Figure S17.** ER reporter architecture and experimental timeline. (a) Domain architecture of the EGFP glycosylation reporter, including the signal sequence, EGFP, linker segments, the N-glycosylation sequon and KDEL. (b) Annotated C-terminal reporter sequence, showing the sequon, FLAG tag and KDEL. (c) Transfection, protein expression, UV decaging, time-course sampling and immunoblot analysis.

**Figure S18**

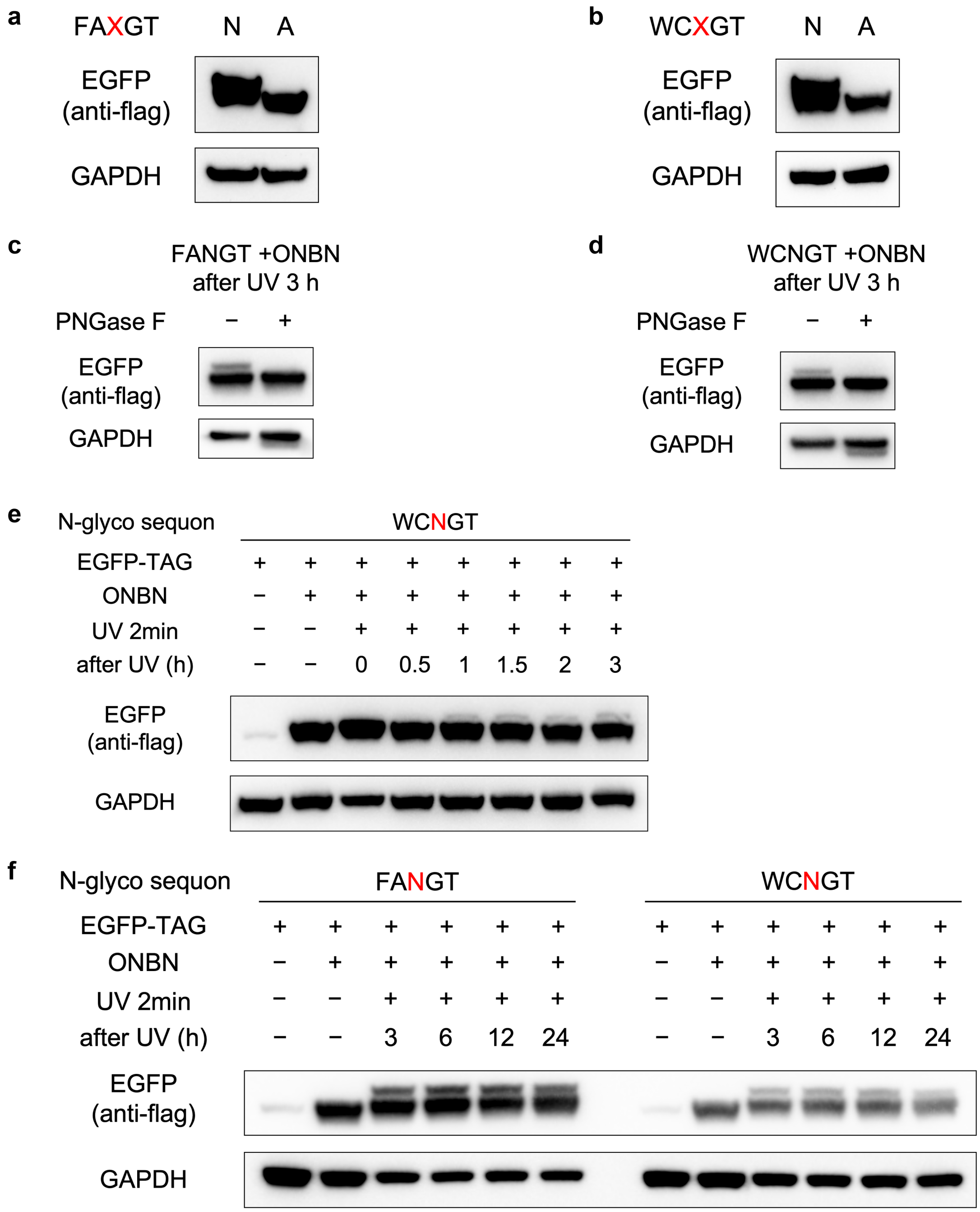


**Figure S18.** Additional glycosylation-reporter validation. (a, b) FANGT- and WCNGT-derived reporters carrying Asn or Ala at the acceptor position. (c, d) PNGase F treatment of the FANGT- and WCNGT-derived reporters collected 3 h after UV irradiation. Anti-Flag detects the EGFP reporter, and GAPDH serves as the loading control. (e) WCNGT reporter series through 3 h. (f) FANGT and WCNGT reporter series at 3, 6, 12 and 24 h. Samples were irradiated for 2 min where indicated. Immunoblots are representative of three independent experiments.

Figure S19

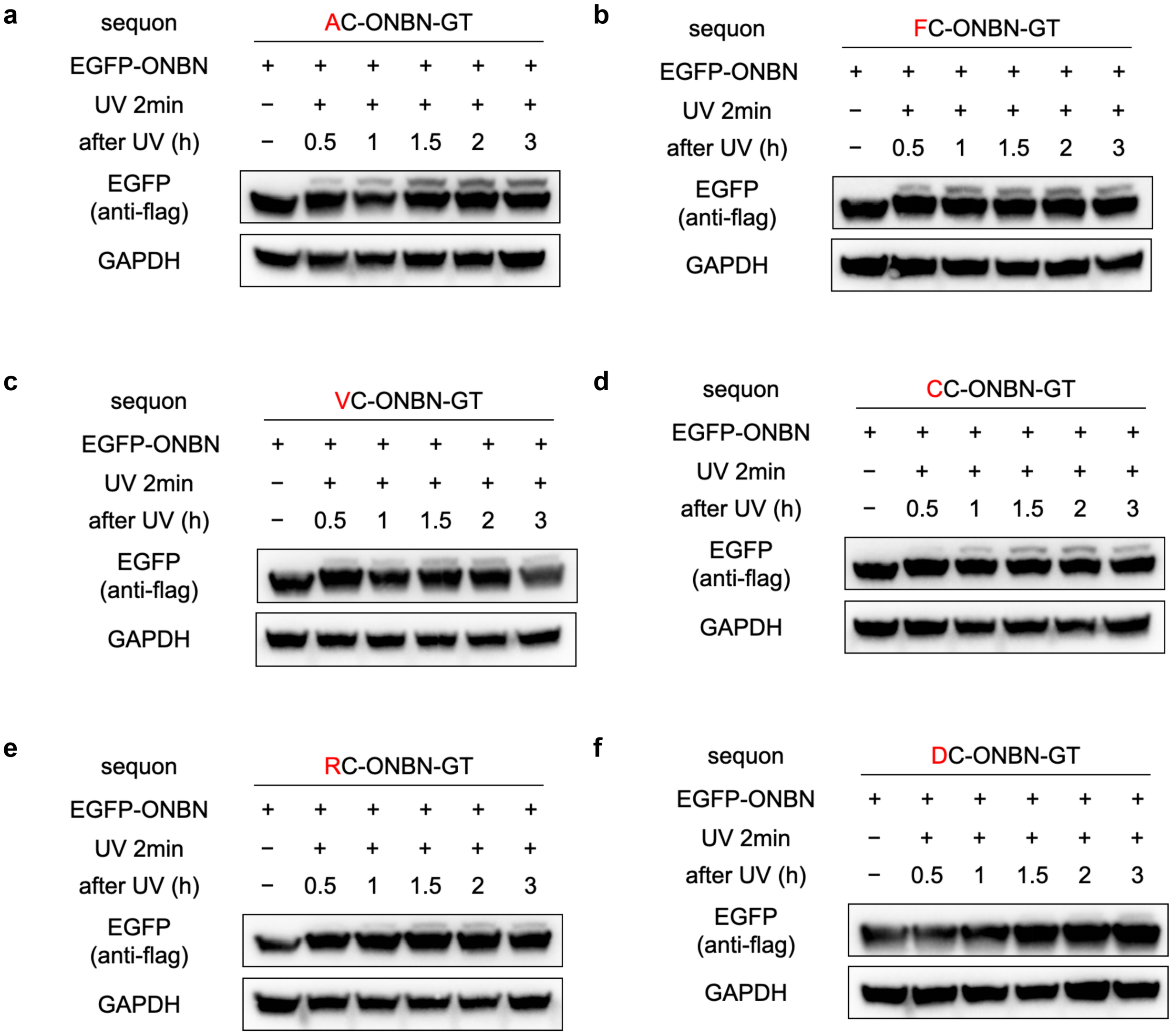


**Figure S19.** Time-resolved glycosylation of the XCNGT reporter series. (a–f) Original anti-Flag immunoblots for X = A, F, V, C, R and D, respectively, with GAPDH as the loading control. The displayed series is 0, 0.5, 1, 1.5, 2 and 3 h. Quantitative source values are in the XCNGT worksheet of ONBN_Source_Data.xlsx. Immunoblots are representative of three independent experiments.

**Figure S19 continued**

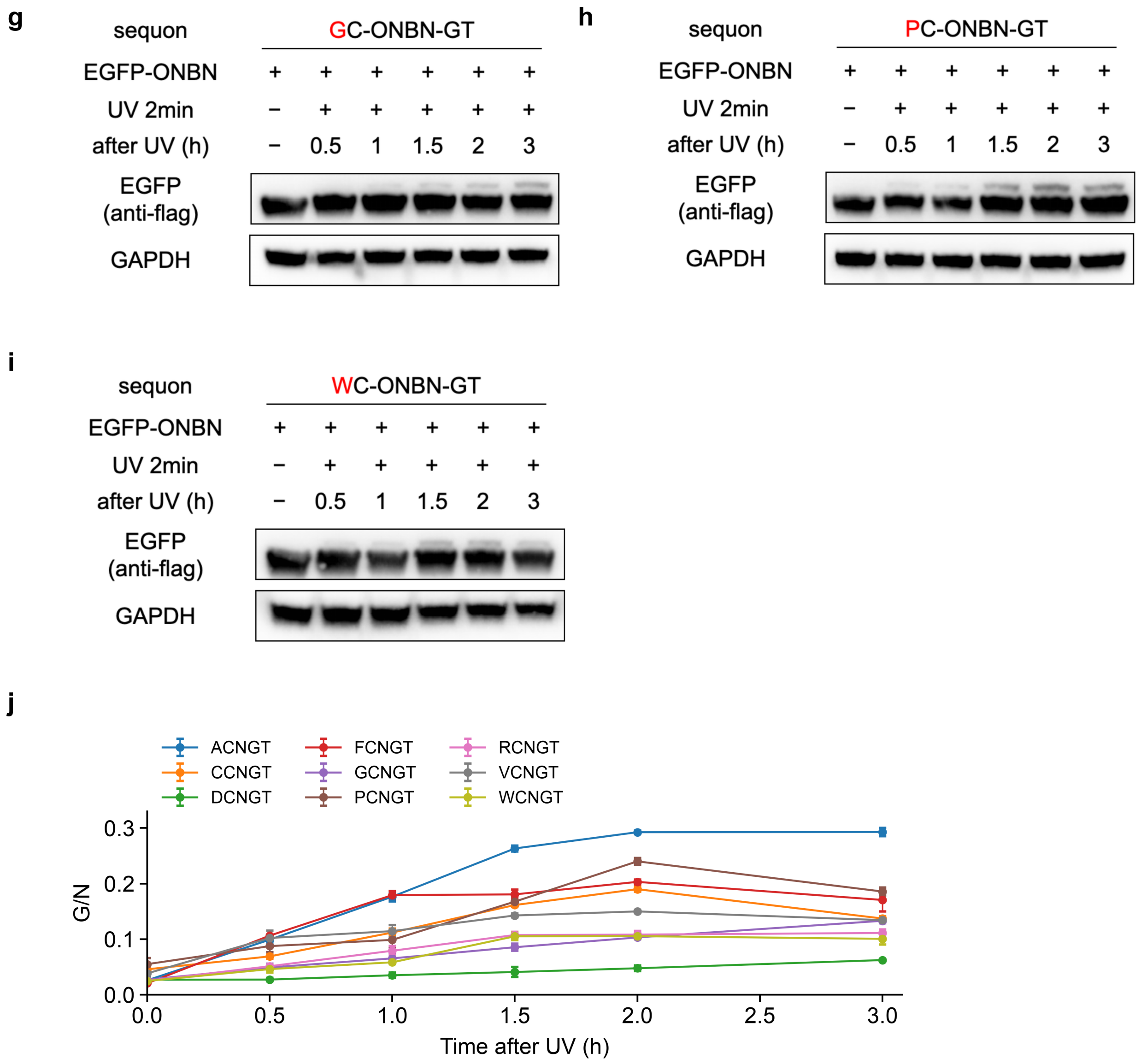


**Figure S19 continued.** (g–i) Corresponding immunoblots for X = G, P and W, respectively. (j) Complete overlaid G/N time courses for all nine variants, shown as mean ± SD ($n$ = 3). G/N is the background-corrected glycosylated-band intensity divided by the nonglycosylated-band intensity. The original G/N values and SD were retained when the plotting frame was resized. Immunoblots are representative of three independent experiments.

#### NMR Spectra

##### Protected ONBN intermediate — $^{1}H$ NMR ($CDCl_3$)

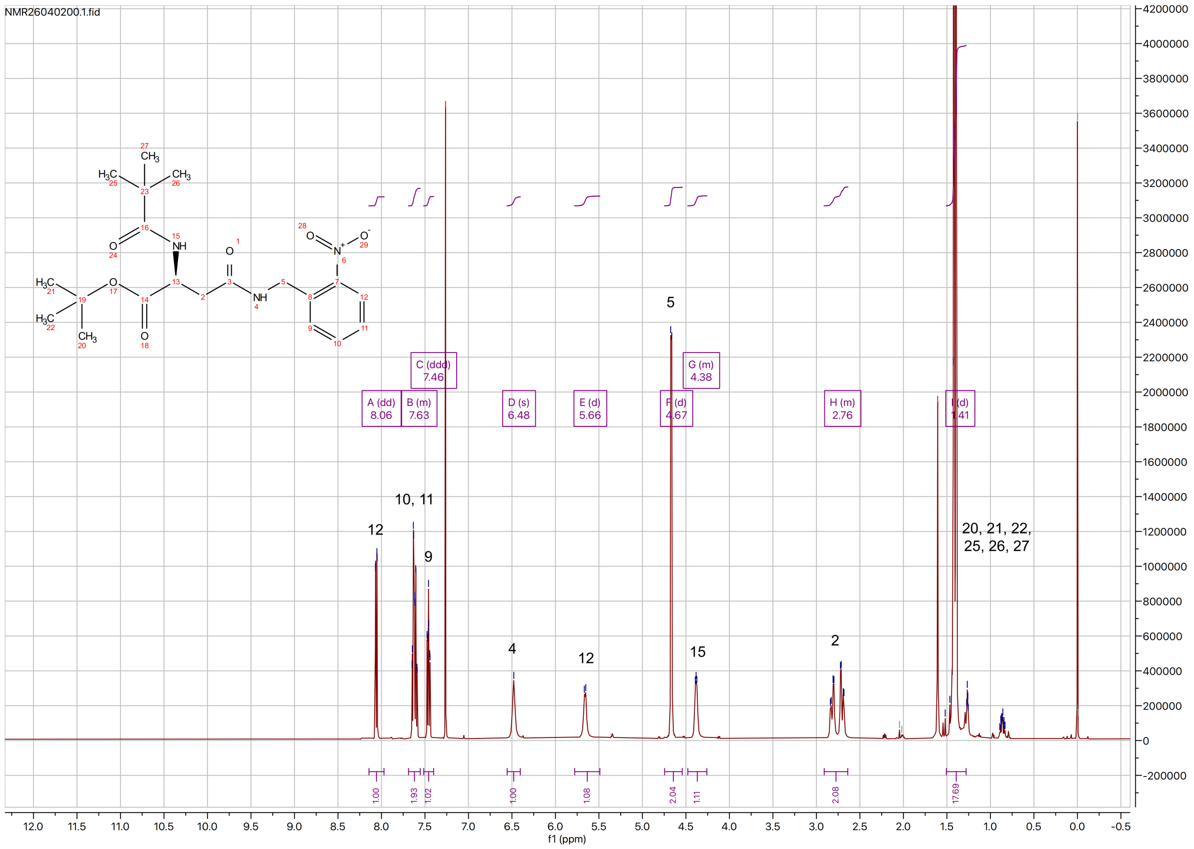


##### Protected ONBN intermediate — $^{13}C$ NMR ($CDCl_3$)

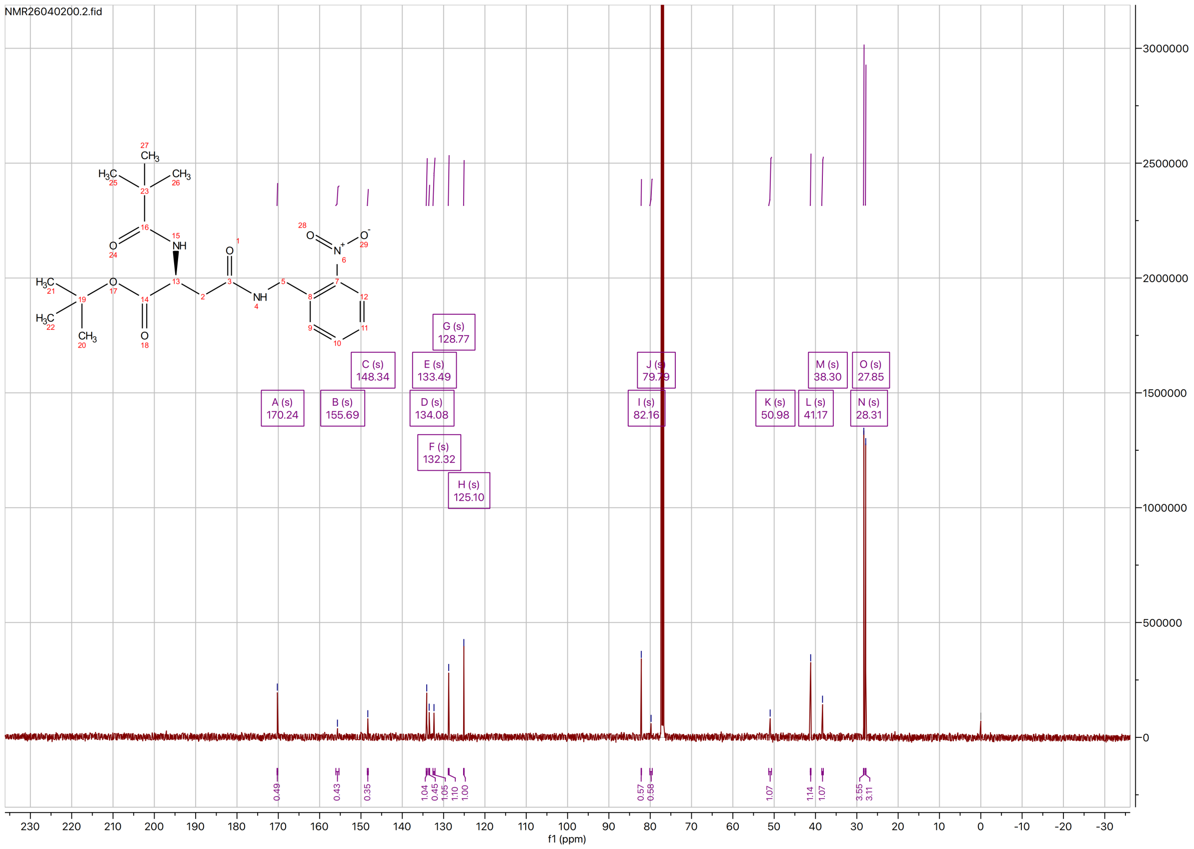

#### ONBN — [1]H NMR (D$_2$O)

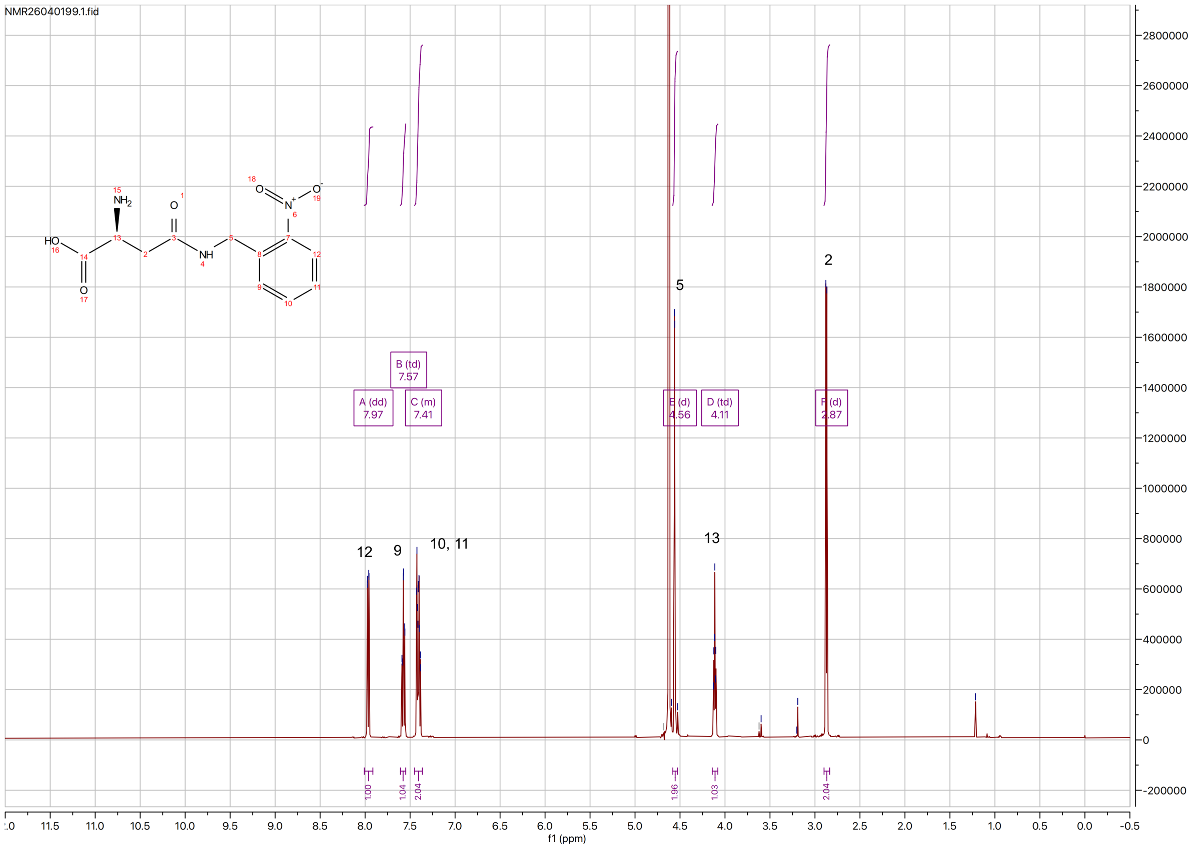


#### ONBN — [13]C NMR (D$_2$O)

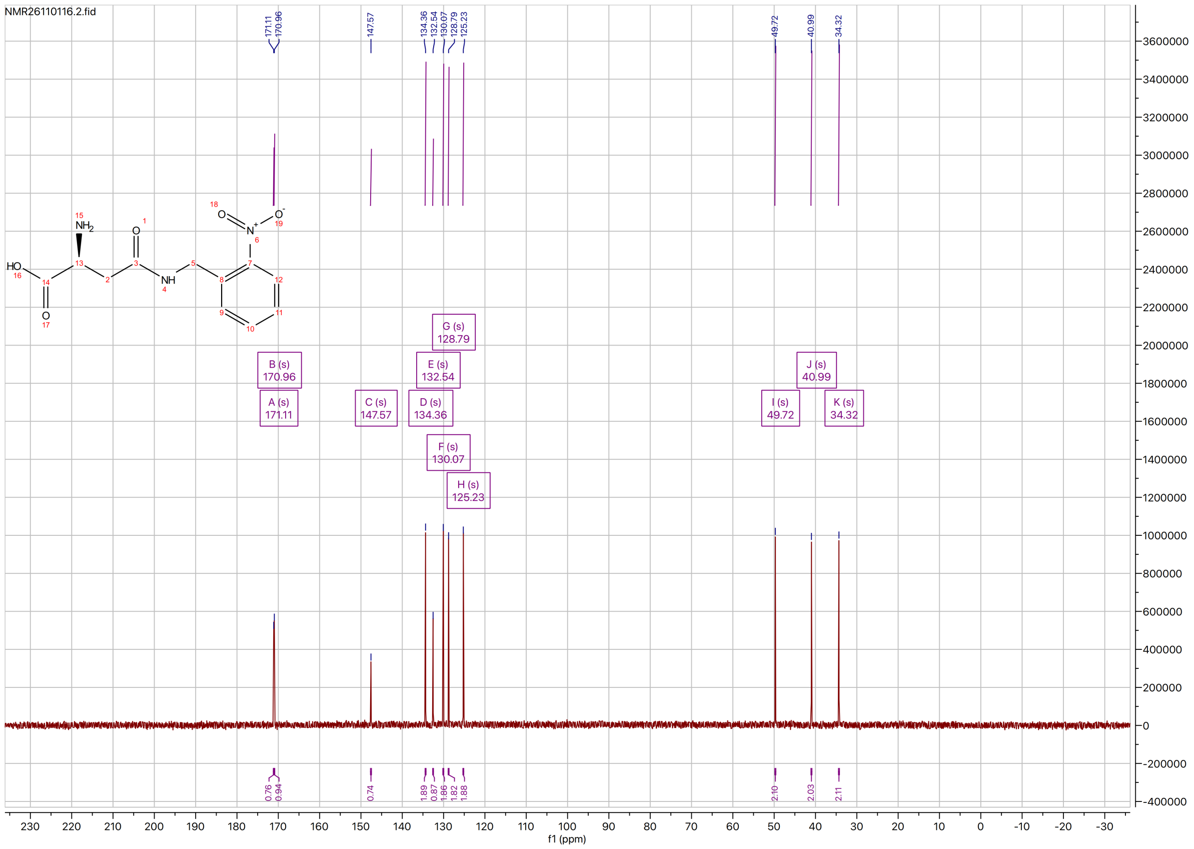

### High Resolution Mass Spectra

#### Protected ONBN intermediate

##### Peking University Mass Spectrometry Sample Analysis Report

**Analysis Info**

| | | | |
|---|---|---|---|
| Analysis Name | FTMS-26040060_000001.d | Acquisition Date | 4/23/2026 8:31:01 PM |
| Sample | Protected-ONBN | Instrument | Bruker Solarix XR FTMS |
| Comment | ESI, MeOH | Operator | Peking University |

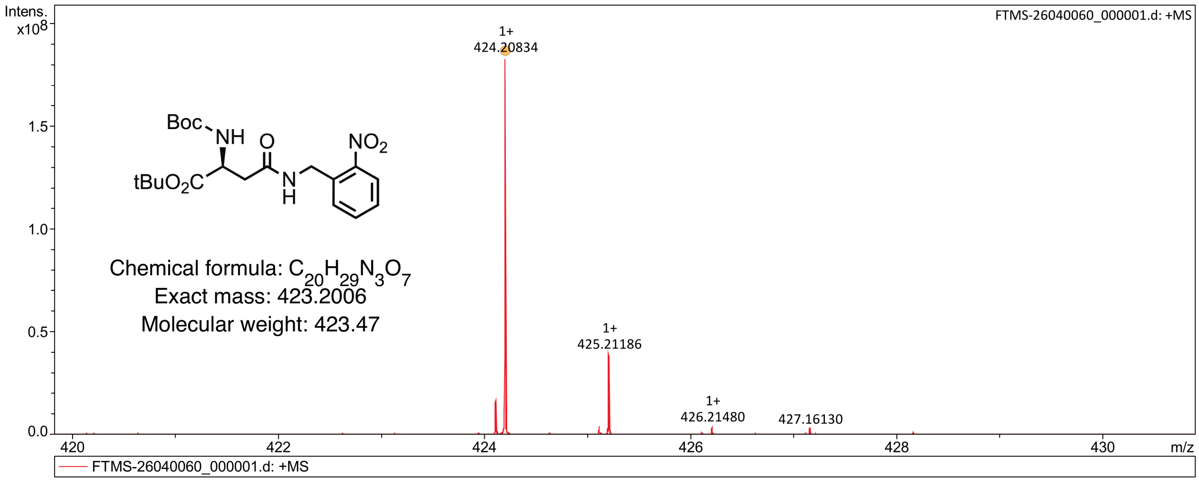


| Meas. m/z | # | Ion Formula | Score | m/z | err [ppm] | Mean err [ppm] | mSigma | rdb | e¯ Conf | N-Rule |
|---|---|---|---|---|---|---|---|---|---|---|
| 424.208341 | 1 | C20H30N3O7 | 100.00 | 424.207827 | -1.2 | -1.9 | 14.5 | 8.0 | even | ok |

#### ONBN

##### Peking University Mass Spectrometry Sample Analysis Report

**Analysis Info**

| | | | |
|---|---|---|---|
| Analysis Name | FTMS-26040061_000001.d | Acquisition Date | 4/23/2026 8:35:16 PM |
| Sample | ONBN | Instrument | Bruker Solarix XR FTMS |
| Comment | ESI, MeOH | Operator | Peking University |

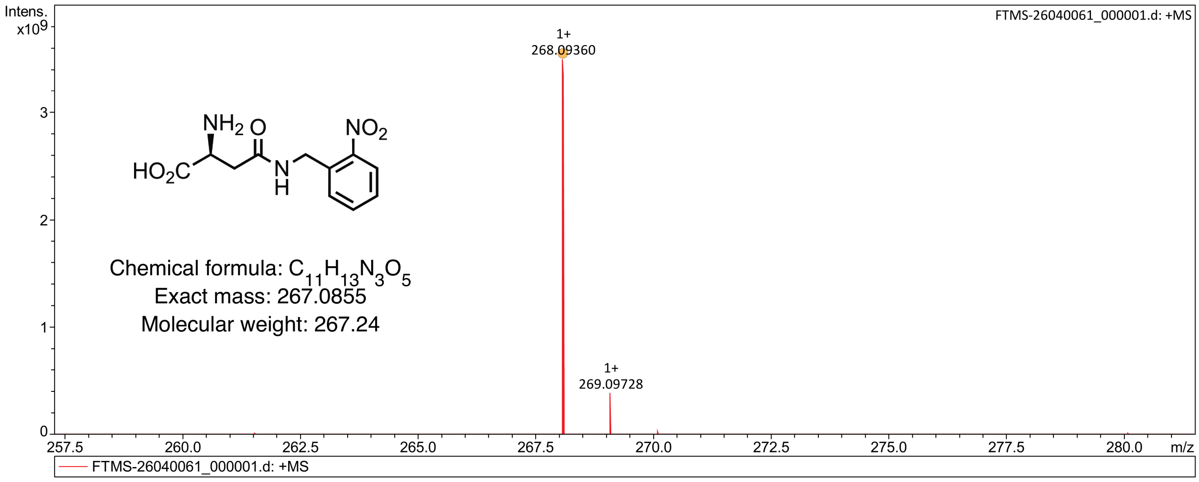


| Meas. m/z | # | Ion Formula | Score | m/z | err [ppm] | Mean err [ppm] | mSigma | rdb | e¯ Conf | N-Rule |
|---|---|---|---|---|---|---|---|---|---|---|
| 268.093602 | 1 | C11H14N3O5 | 100.00 | 268.092797 | -3.0 | -4.4 | 9.4 | 7.0 | even | ok |
